# A temperate endotherm trades thermoregulation for self-preservation: stress-induced changes in surface temperature as thermoregulatory trade-offs

**DOI:** 10.1101/2020.04.06.027623

**Authors:** Joshua K. Robertson, Gabriela F. Mastromonaco, Gary Burness

## Abstract

Coping with stressors can require substantial energetic investment, and when resources are limited, such investment can preclude simultaneous expenditure on other biological processes. Among endotherms, energetic demands of thermoregulation can be immense, yet whether a stress response is sufficient to induce changes in thermoregulatory investment appears unexplored.
We tested the hypothesis that stress-induced changes in surface temperature, a well-documented phenomenon across vertebrates, stem from a trade-off between thermoregulation and stress responsiveness, whereby individuals seek to reduce energetic expenditure on thermoregulation in challenging environments (the “Thermoprotective Hypothesis”). We predicted that surface temperature and dry heat loss of individuals that are resource-limited would fall under stress exposure at low ambient temperatures and rise under stress exposure at high ambient temperatures when compared with non-resource limited individuals.
To test our predictions, we exposed Black-capped Chickadees to rotating stressors and control treatments (n_days/treatment_ = 30; paired treatments) across an ambient temperature gradient whilst remotely monitoring both feeding behaviour and surface temperature.
Supporting the Thermoprotective Hypothesis, our results showed that: 1) social subordinates (*n* = 12), who fed less than social dominants and alone suffered stress-induced declines in mass, displayed significantly larger changes in surface temperature following stress exposure than social dominants (*n* = 8), and 2) stress-induced changes in surface temperature significantly increased heat conservation at low ambient temperature, and heat dissipation at high ambient temperature among social subordinates alone.
These results suggest that Black-capped Chickadees adjust their thermoregulatory strategies under stress when resources are limited and support the hypothesis that stress-induced changes in temperature are functionally significant.

Data Availability Statement: All data and R code used for the construction of this study are available at: https://datadryad.org/stash/share/QzmRV875_S5IYpMT0jxOYIsoO-ua0T3LX6AA9u-XpdQ.

## 1. Introduction

The vertebrate stress response is characterised by coordinated behavioural and physiological modifications that enhance an individual’s ability to cope with perceived challenges, or “stressors”. Over the past century, research pertaining to stressor-induced physiological modifications has undergone overwhelming expansion (e.g. Cannon, 1932; Romero, Dickens, & Cyr, 2009; Selye, 1950; lay literature: Sapolsky, 2004), and this expansion has permitted the identification of numerous and multisystemic adaptations among vertebrates that not only serve to mediate stressors acutely, but also serve to minimize the potential of wear-and-tear acquired from long-term activation of emergency pathways (that is, to reduce allostatic load). To ecologists, adaptations of this latter form are of particular interest as they commonly involve re-allocation of resources among biological processes (e.g. “trade-offs”) that can reveal hierarchies of investment in a species. To date, the vast majority of studies investigating stress-induced trade-offs have focused on changes to reproduction (Calisi, Rizzo, & Bentley, 2008; Grachev, Li, & O’Byrne, 2013; Kinsey-Jones et al., 2009; Kirby, Geraghty, Ubuka, Bentley, & Kaufer, 2008) and immune function (Stier et al., 2009; Svensson, R, Koch, & Hasselquist, 1998), probably owing to the importance of each process with respect to fitness, and the relatively high costs associated with their activation and maintenance (Martin, Scheuerlein, & Wikelski, 2003; Ots, Kerimov, Ivankina, Ilyina, & Hörak, 2001; Speakman, 2008; Vleck, 1981, but see Merlo, Cutrera, Luna, & Zenuto, 2014). Among endothermic vertebrates, however, energetic costs of maintaining a balanced core temperature can often overwhelm those of both reproductive and immune function (Burness, Armstrong, Fee, & Tilman-Schnidel, 2010; King & Swanson, 2013; Nord, Sandell, & Nilsson, 2010), and failing to maintain a balanced core temperature can be equally as detrimental to individual fitness as failing to maintain reproduction or immunity. Nevertheless, trade-offs with respect to thermoregulatory investment in the presence of stressors are largely unexplored (but see Robertson, Mastromonaco, & Burness, 2020).

Changes in body temperature in response to stressors have been widely reported across vertebrate taxa (e.g. squamates: Cabanac & Gosselin, 1993; birds: Greenacre & Lusby, 2004; fish: Rey et al., 2015; old-world primates: Parr & Hopkins, 2000; rodents: Dymond & Fewell, 1998; Yokoi, 1966) and have been documented in medical literature for nearly two thousand years (Galen, 2^nd^ century CE; Yeo, 2005). Although the proximate mechanisms driving thermal responses to stressors have since been well characterized (e.g. haemetic redistribution, changes in brown adipose metabolism, secretion of pyrogenic cytokines; Kataoka, Hioki, Kaneko, & Nakamura, 2014; reviewed in Oka, Oka, & Hori, 2001), an adaptive value of this phenomenon remains contended. Indeed, whilst some studies posit that thermal fluctuations *per se* may endow an individual with immunological advantages during coping (Oka et al., 2001), others argue that stress-induced thermal fluctuations are merely “spandrels” that emerge from haemetic redistribution to mitigate risk of haemorrhage (Bartlett, 1912; Jerem, Herborn, McCafferty, McKeegan, & Nager, 2015; Jerem et al., 2018), or as artifacts of experimental design (Andreasson, Nord, & Nilsson, 2019; Nord & Folkow, 2019).

Recently, Robertson et al. (2020) proposed that an adaptive value of stress-induced changes in body temperature may be understood when contextualised according to an individual’s perceived thermoregulatory costs, and when changes in body temperature are observed at the level of an individual’s surface tissues. Specifically, they hypothesised that changes in surface temperature following stress exposure occur to reduce thermoregulatory costs incurred during activation of a stress response by reducing dry heat loss at temperatures below thermoneutrality, and increasing dry heat loss at temperatures above thermoneutrality. Under this hypothesis (henceforth, the “Thermoprotective Hypothesis”; Robertson et al., 2020), stress-induced changes in surface temperature may be viewed as products of a trade-off, whereby individuals allocate energy away from metabolically expensive thermoregulatory processes (e.g. shivering and non-shivering thermogenesis in cold, and evaporative cooling in heat; costs reviewed in McKechnie et al., 2016), in favour of self-preservation. Supporting the Thermoprotective Hypothesis, numerous studies have reported an influence of ambient temperature on both the magnitude and direction of stress-induced thermal responses at the skin (Jerem, Jenni-Eiermann, McKeegan, McCafferty, & Nager, 2019; Nord & Folkow, 2019; Robertson et al., 2020; Yokoi, 1966). These studies suggest that such changes may indeed be leveraged for thermoregulatory purposes. Additionally, in some studies, both skin temperature and superficial blood volume have been shown to rise following stress exposure (e.g. at the ears of Domestic Rabbits, *Oryctolagus cuniculus*, and wattles of Domestic Chickens, *Gallus gallus*; Herborn, Jerem, Nager, McKeegan, & McCafferty, 2018; Yokoi, 1966) rather than fall. Such trends directly contrast the hypothesis that stress-induced thermal responses are merely functionally neutral corollaries of blood-loss mitigation strategies. Whether stress-induced changes in surface responses vary according to an individual’s energetic or resource availability, and thus represent true trade-offs (reviewed in Bruener and Berk, 2019), however, remains untested.

In this study, we sought to understand whether stress-induced changes in surface temperature emerge from trade-offs between thermoregulation and stress responsiveness, as assumed by the Thermoprotective Hypothesis. As such, we asked: 1) whether an individual’s access to energy (here, both as external food and internal mass) dictates the magnitude of their surface temperature response to a stressor, and 2) whether the direction of an individual’s surface temperature response to stress exposure enhances conservation of energy by decreasing dry heat loss in cold environments, and increasing dry heat dissipation in warm environments. To answer these questions, we exposed Black-capped Chickadees (*Poecile atricapilus*; Linnaeus, 1766), a temperate endothermic species, to randomised and repeated stressors across a naturally occurring temperature gradient, whilst monitoring surface temperature and total rate of dry heat loss (q_Tot_) by infrared thermography. Next, we tested whether resource availability influenced the magnitude of stress-induced surface temperature responses by leveraging naturally occurring, social stratifications in resource access as experimental groupings. Specifically, in Black-capped Chickadee social groups, socially dominant individuals typically have greater access to resources than socially subordinate individuals (supplemental food, mass, and fat deposition; Ficken, Weise, & Popp, 1990; Glase, 1973; Smith, 1991; van Oort, Otter, Fort, & McDonnell, 2007, but see Schubert et al., 2007). Social status (i.e. socially subordinate or socially dominant) may therefore serve as a natural, experimental modulator of resource access. Under the Thermoprotective Hypothesis, we predicted that the surface temperature and q_Tot_ of socially subordinate individuals would be more sensitive to stress exposure than that of socially dominant individuals across an ambient temperature spectrum. More specifically, we predicted that social subordinates (who typically have lower access to available food, and lower fat stores) would experience a greater fall in surface temperature and q_Tot_ at low ambient temperatures, and a greater rise in surface temperature and q_Tot_ at high ambient temperatures than social dominants when exposed to repeated stressors. We predicted this because lower access to resources in subordinate individuals would preclude simultaneous investment in metabolic heating or cooling (i.e. shivering and non-shivering thermogenesis, and evaporative cooling) and self-protection (Fig. 1).

**Figure 1.**
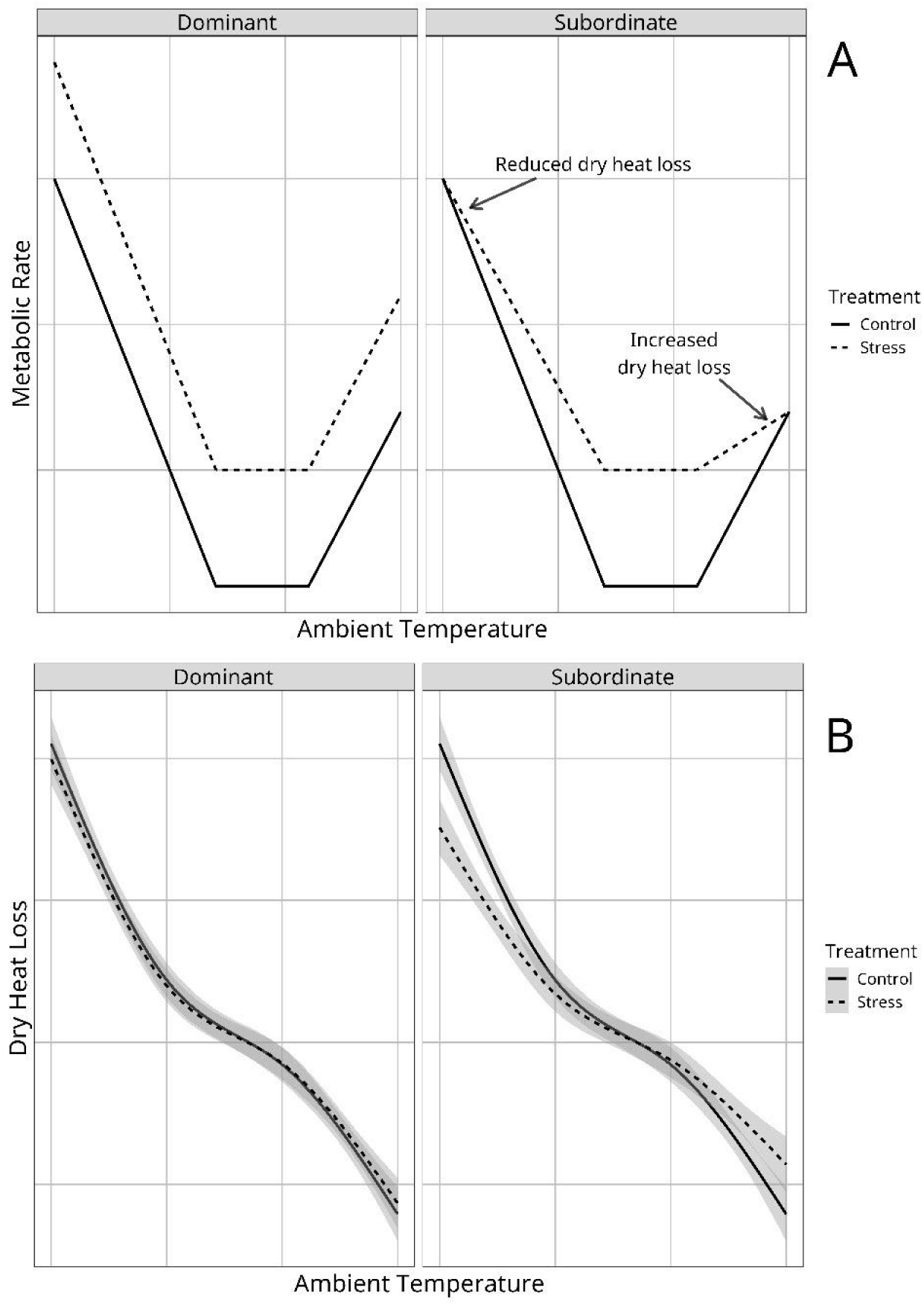
Predicted effects of stress-exposure and resource limitation (dictated by social status) on dry heat transfer at the skin under the Thermoprotective Hypothesis. **A** | Theoretical relationships between ambient temperature and metabolic rate under stress-exposure, and according to energetic availability (stored in food and mass), where social subordinates but not social dominants are energetically limited. Socially dominant individuals are predicted to increase their metabolic rate in response to stressors regardless of ambient temperature (within a range observable in our study). Social subordinates are predicted to relax energetic expenditure toward thermoregulation in favour of stress responsiveness at temperatures outside of thermoneutrality. Solid lines represent theoretical metabolic responses to control conditions, dashed lines represent theoretical responses to stress exposure. **B** | Predicted effects of stress exposure on dry heat loss across ambient temperature, for socially subordinate and socially dominant individuals. Socially dominant individuals are not predicted to change rates of dry heat loss following exposure to stressors. Contrastingly, social subordinates are predicted to reduce their dry heat loss under stress exposure in the cold, and increase their dry heat loss under stress exposure in the heat. Solid lines represent theoretical metabolic responses to control conditions, dashed lines represent predicted theoretical responses to stress exposure.

To our knowledge, this is the first study to test whether thermal response to stress exposure might represent functional reallocation of resources to accommodate heightened demands experienced during coping. Furthermore, it is also one of few to explore the presence of a trade-off under stress exposure with respect to ecologically relevant resource constraints (i.e. those induced by social hierarchies; Aghajani et al., 2013; Proctor, Freeman, & Brown, 2010).

## 2. Materials and Methods

All protocols used for animal capture, handling, and experimental manipulation were approved by the Government of Canada (Environment and Climate Change Canada; permit #10756E) and by the Trent University Animal Care Committee (AUP #24614).

### 2.1 Capture and sampling of Black-capped Chickadees

Black-capped Chickadees used for experimentation (n = 20) were captured within a 100 km^2^ region of Southern Ontario (Canada), during March and April of 2018. We divided capture efforts across both urban and rural locations (n = 3 sites per category) to control for possible effects of urbanisation on stress-induced changes in surface temperature alone (Abolins-Abols, Hope, & Ketterson, 2016). Urban capture locations included the cities of Cambridge (43.379°N, 80.353°W), Guelph (43.330°N, 80.150°W), and Brantford (43.135°N, 80.344°W), whilst rural capture locations included the greater townships of Erin and Corwhin (43.762°N, 80.153°W and 43.509°N, 80.090°W respectively), and the Ruthven Park National Historic Site (42.980°N, 79.875°W). Our final sample population included 10 individuals captured from urban locations (n_female_ = 5, n_male_ = 5), and 10 individuals captured from rural locations (n_female_ = 5, n_male_ = 5).

All Chickadees were captured using modified potter traps following Robertson et al. (2020). Immediately following capture, Chickadees were blood sampled for genetic sexing by brachial venipuncture (approximately 50 μ L), then assigned a unique combination of one government-issued stainless steel leg band (size 0; Environment and Climate Change Canada; Bird Banding Office) and two coloured leg bands (Darvic, 2.3 mm interior diameter; Avinet, Portland, Maine, USA). Birds were then weighed, measured (mass: nearest 0.1 g using a digital platform scale; left tarsus and flattened wing-chord; nearest mm using analogue calipers), and secured in transportation cages (30.0 cm x 30.0 cm x 15.0 cm; l x w x h) for translocation to long-term holding facilities at the Ruthven Park National Historic Site, Cayuga, Ontario, Canada (< 90 km; 2 hours travel by vehicle). Erythrocytes were isolated from blood samples by centrifugation in the field (12,000 rpm), preserved in Queen’s Lysis Buffer (Seutin, White, & Boag, 1991), then stored at 4°C until required for analysis.

### 2.2 Captive maintenance

Housing of Black-capped Chickadees is described elsewhere (Robertson et al., 2020). Briefly, all birds were randomly allocated to one of four visually-isolated flight enclosures (n = 5 per flight enclosure; 1.83 cm x 1.22 m x 2.44 m; l x w x h), each supplied with a roosting box (60 cm x 20 × 20 cm; l x w x h), one roosting tree (White Cedar, *Thuja occidentalis*; 1.0 m), and two perching branches (approximately 80 cm in length). Enclosures were fitted with one feeding platform (4 dm^2^) placed at approximately 1.2 m in height, that could be provisioned from behind an opaque barrier to ensure that birds were blind to the presence of experimenters. Additionally, a small, water-tight camera-box perforated with two small holes (30 mm in diameter) was secured to an outside wall of each flight enclosure, and positioned parallel to each feeding platform to permit filming of individuals through the perforations during feeding (described in “Thermographic Filming”). Camera-boxes were not visible to Chickadees within flight enclosures and were solely accessed from the exterior of the enclosures.

Chickadees were acclimated to flight-enclosures for a minimum of two weeks before experimentation. During both acclimation and experimentation periods, all birds were fed a mixture of mealworms (*Tenebrio molitor*), crickets (*Acheta domesticus*), sunflower seeds, safflower seeds, shelled peanuts, apple pieces, boiled egg, and Mazuri™ (St Louis, Missouri, USA) Small Bird Maintenance diet *ad libitum*, and were supplied with *ad libitum* fresh water each day.

### 2.3 Experimental stress exposure

To test whether social status influenced the magnitude of stress-induced changes in surface temperature among Chickadees, we employed a rotational stress exposure protocol similar to that used in Rich & Romero (2005). A rotational approach to stress exposure was chosen to circumvent habituation to each individual stressor across the duration of our experiment. To increase statistical power, we used a repeated sampling approach whereby all individuals received both a control and stress exposure treatment, with each treatment persisting for 30 days and being separated by a rest period of two days (total experimental duration of 62 days). To control for possible effects of treatment order, individuals were divided into two groups of ten individuals (n_flight enclosures per group_ = 2; one west facing, and one east facing per grouping), with one group receiving a control treatment followed by a stress exposure treatment, and the other group simultaneously receiving a stress exposure treatment followed by a control treatment.

In stress exposure treatments, individuals were exposed to five or six randomly selected, passive stressors each day, with each exposure persisting for 20 minutes and being separated from previous and subsequently stressors by 1 hour. Possible stressors included; 1) presence of an experimenter within the flight enclosure of target individuals, 2) capture and restraint in an opaque, fabric bag, 3) presence of a mock predator (adult Cooper’s Hawk, *Accipiter cooperii*), 4) presence of a taxidermised conspecific, mounted to the feeding platform of target individuals, 5) presence of a novel object (garden gnome) placed near the feeding platform of target individuals, and 6) complete enwrapment of the flight enclosure with an opaque fabric sheet. Auditory stressors were not used in this experiment to avoid evocation of stress responses in Chickadees housed in nearby flight enclosures. The effectiveness of our rotational stress exposure protocol has been confirmed for Black-capped Chickadees elsewhere (Robertson et al, 2020).

In control treatments, individuals were maintained according to acclimation conditions, and were not handled or exposed to experimenters. Chickadees were returned to their location of capture and released.

### 2.4 Thermographic imaging, digital filming, and environmental data collection

To monitor the surface temperature of Chickadees during experimental treatments, we passively imaged individuals during feeding using a remotely-operated infrared thermographic camera (VueProR™, FLIR, Wilsonville, Oregon, USA; 13 mm lens, 336 × 256 resolution) that was by placed in a water-tight camera-box and oriented toward a focal feeding platform at a distance of 0.5 m (camera-boxes described in “Captive Maintenance”). Thermographic filming was conducted at a frequency of 1 image per second for a minimum of 1 hour per enclosure, per day. Complete thermography protocols are described in Supplemental Methods. Identity of individuals within thermographic images were determined from digital video captured in parallel to thermographic images (Supplemental Methods). All thermographic images and digital video used in this study were captured between 08:00 and 16:00 across a total of 60 days.

Both ambient temperature and relative humidity are known to influence estimates of an object’s surface temperature by infrared thermography (reviewed in Minkina & Dudzik, 2009; Tattersall, 2016). To support accurate calculation of Black-capped Chickadee surface temperature measurements from thermographic images, we therefore collected ambient temperature and relative humidity measurements across the duration of experimentation. Ambient temperature measurements were recorded within flight enclosures during thermographic filming intervals using a ThermoChron iButton™ (1 reading/5 minutes at a resolution of 0.5°C; Maxim Integrated, DS1922L-F5; San Jose, California, USA) placed in the shade, whilst relative humidity readings were collected from a governmental climate repository (Department of Environment and Climate Change Canada; http://climate.weather.gc.ca/; Hamilton A; 22 km from the site of experimentation). Humidity readings were collected at a frequency of 1 reading/hour (the highest resolution available).

### 2.5 Social status determination

Social status of individual Chickadees was estimated from enumerated presence and absence data derived from digital video observations made in parallel to thermographic filming (Supplemental Methods). Here, individual social status was assessed per flight enclosure, using a similar approach to that recommended by Evans et al. (2018). Specifically, presence and absence data at feeding platforms were used to infer displacement events (where the arrival of individual A resulted in the departure of individual B), waiting events (where the arrival of individual B was contingent upon the departure of individual A), and non-agonistic events (where individuals A and B fed simultaneously) among Chickadees, across the duration of our experiment (n_days_ = 60). Individuals involved in displacement or waiting events were then assigned a “win” or “loss” according to whether they received priority access to the feeding platform. Those involved in non-agonistic interactions were assigned “draws”. Total wins, losses, and draws per individual were then used to calculate an index of relative wins in social interactions, and this index was used to infer social status values of Chickadees within each flight enclosure. For this study, we chose to use a randomised Elo rating (Elo, 1978; calculated from randomly sampled days of behavioural observation, with sampling iterated 1000 times; Supplemental Methods) as our index of relative wins because it is thought to outperform other widely used status indices (e.g. David’s Score, ADAGIO, IS&S) when predicting social status in the presence of shallow dominance hierarchies (Sánchez-Tójar, Schroeder, & Farine, 2018).

To assess our confidence in social status estimates, we calculated a “status-stability index” (“SSI”) per individual, which describes the proportion of final status estimates (that is, those assumed upon completion of each random sampling of observation dates) that were found to be equivalent to an individual’s modal social status estimate (similar to van Hooff and Wensing’s directional consistency index [1987], but expanded for application to randomised data). SSI per individual was calculated using the following equation;

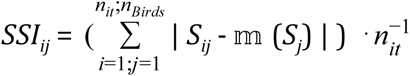

where, *S* represents the estimated social status of an individual *j* according to their Elo rating that was calculated following a randomised iteration *i*, 𝕞 (*S*_*j*_) represents the modal social status of individual *j* across all iterations, *i* (consequently, the final estimated social status for an individual *j*), *n*_*it*_ represents the total number of randomised iterations, and *n*_*Birds*_ represents the total number of individuals for which social status were estimated.

### 2.6 Calculation of surface temperature and heat transfer from thermographic images

Surface temperature measurements (maximum temperature of the periorbital region, or “eye region”) of Black-capped Chickadees were derived from thermal images in R (approximately 230 pixels; R version 3.6.1; Core Team, 2019) according to previously described methods (Robertson et al., 2020). Emissivity of Black-capped Chickadee integument was estimated as 0.95, according to Best & Fowler (1981). We chose to assess surface temperature at the eye region alone because it is readily imaged during feeding events, it contains exposed integument that is uninsulated by keratinous tissues (ie. leg scales or feathers; discussed by Jerem et al., 2018), and it has previously been shown to be thermally responsive to stress exposure (e.g. Jerem et al., 2015; Robertson et al., 2020). Additionally, we chose to monitor the maximum of the eye region temperature rather than the mean of the eye region temperature because it is thought to be less vulnerable to measurement error, and more consistent across image angles (Jerem et al., 2015, 2018). Similar to Robertson et al. (2020), surface temperature measurements were only extracted from images where identity of the focal individual could be determined by video observations, and when the focal individual was not in motion (to avoid underestimation of surface temperature; discussed in Jerem et al., 2018). In total, data from 6431 thermal images were used for this study, with 3034 images captured during control treatments (social subordinates: n_images_ = 1494, social dominants: n_images_ = 1540) and 3397 images during stress exposure (social subordinates: n_images_ = 1943, social dominants: n_images_ = 1454).

We quantified the rate of heat transfer (mW) from Chickadees using surface temperature measurements (derived from thermographic images) and equations described by Ward et al. (1999), McCafferty et al. (2011) and Nord & Nilsson, (2019) (summarised in Robertson et al., 2020; further details in Supplemental Methods). Heat transfer rates were therefore limited to those experienced at the eye region alone. Because our Chickadees were housed in semi-protective flight enclosures that provided shelter from wind, and because conductive heat transfer from the eye region to a medium other than air was unlikely, we assumed that *q*_*Tot*_ was equal to the sum of convective and radiative heat transfer alone (*q*_*Conv*_ and *q*_*Rad*_ respectively), at a given second. Eye region surfaces were treated as planar, ovoid structures with a vertical diameter of 1.0 cm and a horizontal diameter of and 1.1 cm (surface area = 0.864 cm^2^). All heat transfer calculations were conducted in R and final *q*^*Tot*^ values were multiplied by two to represent total heat transfer across both eye regions of an individual.

### 2.7 Sex determination

In Black-capped Chickadees, males typically hold higher social ranks than females (Glase, 1973; Smith, 1976). To ensure that effects of sex alone did not bias our analyses, we determined the sex of all experimental individuals according to methods described by Griffiths et al. (1996) and Fridolfsson & Ellegren (1999). To do so, we isolated whole genomic DNA from erythrocyte-lysis samples by phenol:chloroform:isoamyl (25:24:1) extraction and 2-propanol precipitation, then amplified intron 16 of the chromohelicase DNA-binding gene (or “CHD” gene) by PCR. Intron 16 of the CHD gene differs in base-pair length between copies located on the W and Z chromosomes. The sex of individuals was therefore determined by size-separation of PCR amplicons on 3% agarose gels (120 V).

### 2.8 Statistical analyses

All statistical analyses were conducted in R, and α levels were set to 0.05. All generalized additive mixed-effects models (“GAMMs”) were fit with restricted maximum likelihood (“REML”) in the R package “mgcv” (Wood, 2011), as recommended by Simpson (2018).

### 2.8.1 Effect of social status on stress-induced changes in surface temperature

We predicted that subordinate individuals – who have reduced access to supplied food (*p* = 0.005; Supplemental Results; SFig. 1), and experience loss of mass during stress exposure treatments (*p* = 0.023; Supplemental Results; SFig. 2) – would elicit a more robust change in surface temperature following stress exposure than dominant individuals. To test whether social status influenced the magnitude of thermal responses to stress exposure across ambient temperature, we used a GAMM with maximum eye region temperature (mean °C for a given hour of observation; n_observations_ = 1027 across n = 60 days) as the response variable (Gaussian distributed). Use of an additive model (in place of a linear model) was chosen to best capture the non-linear relationship between surface temperature and ambient temperature that was previously reported in Black-capped Chickadees (Robertson et al., 2020). Treatment (binomial factor; “control” or “stress-induced”), social status (binomial factor; “dominant” or “subordinate”), and sex (binomial factor) were each included as linear predictors in our model, and an interaction between treatment and social status was also included to account for differential responses to stress exposure across social hierarchies that may occur independently of ambient temperature. An effect of ambient temperature on the surface temperature of Chickadees was tested by inclusion of mean hourly temperature (°C) as a cubic regression spline, with four knots to capture non-linearity and circumvent model over-fit. To test whether the effect of treatment on surface temperature varied across ambient temperature (previously reported in Robertson et al., 2020), we included mean hourly temperature and treatment as a tensor product predictor, with treatment being represented as a smooth factor and mean hourly temperature again being represented as a cubic regression spline with four knots. Importantly, to test the effect of social status on the magnitude of an individual’s thermal response to stress across ambient temperature, a linear interaction between social status, and our hourly temperature and treatment tensor product was included with first-order derivative penalisation.

**Figure 2.**
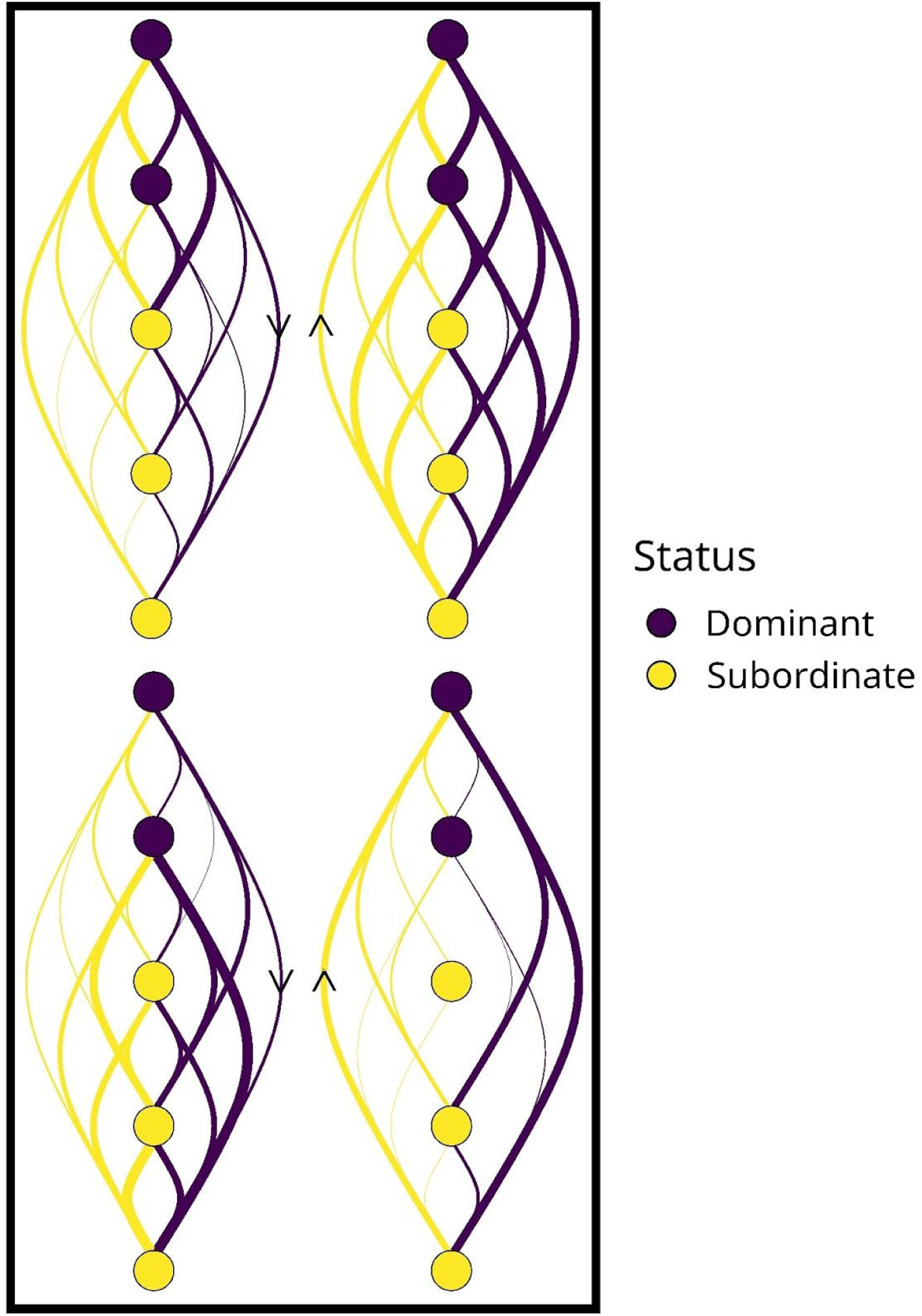
Linear social hierarchies of Black-capped Chickadees (n = 20) across four flight enclosures (n = 5 birds per enclosure) and 60 days of observation. Purple circles represent individuals categorised as socially dominant, whilst yellow circles represent those categorised as socially subordinate; linear order of circles represents that estimated order within a dominance hierarchy (with the most socially dominant individual at the top, and least socially dominant individual at the bottom, as estimated using randomised Elo ratings). Lines represent agonistic interactions between individuals (i.e. supplantations and waiting events), with purple lines representing top-down interactions (dominant to subordinate), and yellow lines representing bottom up interactions (subordinate to dominant). Line width is proportional to the number of interactions between individuals in a dyad.

Because Black-capped Chickadees may display diel changes in body temperature that occur independently of ambient temperature (Robertson et al., 2020), we sought to include time of day (hour) as a predictor in our model, however, we also recognised that the effect of time of day was likely to differ according to flight enclosure orientation (i.e. west- or east-facing), owing to differences in exposure to solar radiation at each hour. For this reason, a tensor product between time of day and flight enclosure orientation (i.e. “east” or “west”; n = 2 per cardinal direction) was included as a predictor in our model, and because stress exposure was previously reported to shift the relationship between time of day and surface temperature, a linear interaction between treatment and our time of day by enclosure tensor product was also included (with first-order derivative penalisation, and with spline conditions as described above). Here, time of day was modeled using a penalised cubic regression spline with 4 knots (cubic shrinkage spline to minimise function size), while enclosure orientation was modeled as a smooth factor. To capture variance explained by date of imaging, individual identity and flight enclosure, random intercepts were included for each factor. Temporal autocorrelation was not detected within individuals in our model (ρ = −0.151).

Changes in surface temperature are likely to yield the greatest energetic consequences at ambient temperatures above or below thermoneutrality. We therefore tested whether the surface temperature of socially dominant and socially subordinate individuals differed between control and stress exposure treatments at ambient temperatures below, at, and above the thermoneutral zone for Black-capped Chickadees (14°C to 30°C; lower limit; Grossman & West, 1977; upper limit; Rising & Hudson, 1974). To do so, we conducted planned comparisons between treatment types within each temperature zone (i.e. sub-thermoneutral, thermoneutral, supra-thermoneutral; n_contrasts_ = 3), and for each level of social status in the R package “emmeans” (Lenth, Singmann, & Love, 2018). Degrees of freedom for each planned comparison were estimated using Kenward-Roger approximations. For this analysis, a broad thermoneutral zone was assumed because our experiment spanned from late winter to early summer, during which the thermoneutral zone of Chickadees is thought to change (Rising & Hudson, 1974).

### 2.8.2 Differences in heat transfer across social status

To understand whether differences in the magnitude of stress-induced changes in surface temperature among individuals impact energy conservation, we asked: 1) whether stress exposure influenced q_Tot_ among individuals, and 2) whether an effect of stress exposure on q_Tot_ was conditional upon the social status of an individual. Because heat transfer from an object is proportional to its surface temperature (*q*_*Tot*_ ∝ *Ts*), we modeled the relationship between *q*_*Tot*_, social status, treatment, and ambient temperature using the same statistical approach as that used for modeling surface temperature in our Chickadees (see “Effect of social status on stress-induced changes in surface temperature”; all predictors remain equivalent). Here, however, maximum eye region temperature was replaced with *q*_*Tot*_ as the Gaussian distributed response variable (n_observations_ = 1027 across n = 60 days).

## 3. Results

### 3.1 Social hierarchies are stable in Black-capped Chickadees

Throughout our experiment, we detected 1006 social interactions between individual Chickadees (mean per individual ± s.d. = 50.300 ± 23.414). Estimates of social status derived from the outcomes of social interactions were stable when calculated
across randomly sampled days of observation (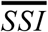 ± s.d. = 78.020%; ± 15.360%; Fig. 2), and stability did not significantly differ among flight enclosures (*F* = 0.122, *d*.*f*. = 3, *p* = 0.946). Stability indices exceeded 50% across all individuals (range = 0.505 - 1.000). Interestingly, social dominance did not appear to be dependent on sex in our study; rather, both males and females were equally represented in socially dominant and socially subordinate positions (socially dominant; n_male_ = 4, n_female_ = 4; socially subordinate; n_male_ = 6, n_female_ = 6).

Unsurprisingly, social subordinates accessed supplemental food less frequently than social dominants (mean feeding visits/hour ± s.e.m.: social subordinates = 5.65 ± 0.626, social dominants = 11.30 ± 1.983; *p* = 0.005; Supplemental Results; SFig. 1), and alone experienced a loss of mass during stress exposure treatments (mean change in mass [Δg] ± s.e.m.: social subordinates = −0.388 ± 0.107, social dominants = +0.032 ± 0.834; *p* = 0.023; Supplemental Results; SFig. 2).

### 3.2 Effects of stress exposure on surface temperature differ according to social status

Maximum eye temperature was positively correlated with ambient temperature in our study animals (*p* < 0.001; Tab.1; Fig. 3a). As predicted by the Thermoprotective Hypothesis, this relationship was significantly influenced by treatment in socially subordinate individuals (social status : tensor product [⊗] of treatment and ambient temperature: *p* = 0.026; Tab.1; Fig. 3a), but not socially dominant individuals (treatment ⊗ ambient temperature: *p* = 0.285; Tab. 1; Fig. 3a). At ambient temperatures below thermoneutrality (< 14°C), the surface temperatures of social subordinates were significantly lower during stress exposure treatments than during control treatments (mean Δ ± s.e.m. = -1.579°C ± 0.508, *t*_*1006*_ = 3.109, *p* = 0.002; Fig. 3b), whilst those of social dominants did not significantly differ between treatment types (mean Δ ± s.e.m, = −0.412°C ± 0.432, *t*_*1006*_ = 0.955, *p* = 0.340; Fig. 3b). Similar trends were observed at ambient temperatures above thermoneutrality (> 30°C), with surface temperatures of social subordinates, but not social dominants, significantly rising under stress exposure treatments (social subordinates: mean Δ ± s.e.m. = 1.220°C ± 0.501, *t*_*1006*_ = 2.436, *p* = 0.015; social dominants; mean Δ ± s.e.m. = 0.636°C ± 0.502, *t*_*1006*_ = 1.267, *p* = 0.205; Fig. 3b). At ambient temperatures within the thermoneutral zone, however, neither the surface temperatures of socially subordinate nor socially dominant individuals significantly differed between treatment types (social subordinates: mean Δ ± s.e.m. = 0.086°C ± s.e.m = 0.193, *t*_*1006*_ = 0.444, *p* = 0.657; social dominants: mean Δ ± s.e.m. = −0.020°C ± 0.210, *t*_*1006*_ = 0.098, *p* = 0.922; Fig. 3b).

**Table 1.** Results of a GAMM testing the influence of social status on the relationship between ambient temperature and surface temperature of Black-capped Chickadees (n = 20) across control and stress-exposed conditions. Mean surface temperature of Chickadees (n_dominant_ = 8; n_subordinate_ = 12) was measured across 60 days (n_days_ = 30 per treatment type), and averaged by hour. Coefficient estimates (β for linear predictors, and mean β, 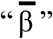, for smooth predictors) and standard errors around coefficient estimates (s.e.m.) are presented for each fixed-effect predictor. Degrees of freedom (*d*.*f*.) are reported for linear predictors, and estimated degrees of freedom (*e*.*d*.*f*.) for smooth predictors. Asterisks (*) signify a significant effect at α = 0.05, and ⊗ represents a tensor product; social dominants: n_measurements_ = 458; social subordinates: n_measurements_ = 569. This model explained 74.941% of deviance in surface temperature data.

| Linear Predictors |  |  |  |  |  |
| --- | --- | --- | --- | --- | --- |
| Predictor | Coefficient ($\beta$) | s.e.m. | *d.f.* | *t* | *p* |
| Intercept | 33.943 | 0.541 | 1 | 62.735 | <0.001* |
| Status | 0.119 | 0.315 | 1 | 0.377 | 0.706 |
| Sex | -0.047 | 0.181 | 1 | -0.260 | 0.795 |
| Treatment | 0.036 | 0.273 | 1 | 0.132 | 0.895 |
| Treatment:Status | 0.044 | 0.125 | 1 | 0.351 | 0.726 |
| Smooth Predictors |  |  |  |  |  |
| Predictor | Mean Coefficient ($\bar{\beta}$) | Mean s.e.m. | *e.d.f.* | *F* | *p* |
| Ambient Temperature | 6.980 | 0.458 | 2.447 | 157.261 | <0.001* |
| Ambient Temperature $\otimes$ Treatment | -0.405 | 0.600 | 2.618 | 0.802 | 0.285 |
| Status: Ambient Temperature $\otimes$ Treatment | -0.217 | 0.480 | 2.626 | 1.154 | 0.026* |
| Time of Day $\otimes$ Enclosure Orientation | -0.018 | 0.581 | 5.429 | 7.039 | <0.001* |
| Treatment : Time of Day $\otimes$ Enclosure Orientation | -0.103 | 0.542 | 3.287 | 4.613 | <0.001* |
| Random Effects |  |  |  |  |  |
| Predictor | Variance |  |  |  |  |
| Bird Identity | 0.471 |  |  |  |  |
| Date of Photo | 0.818 |  |  |  |  |
| Enclosure Identity | 0.818 |  |  |  |  |

**Figure 3.**
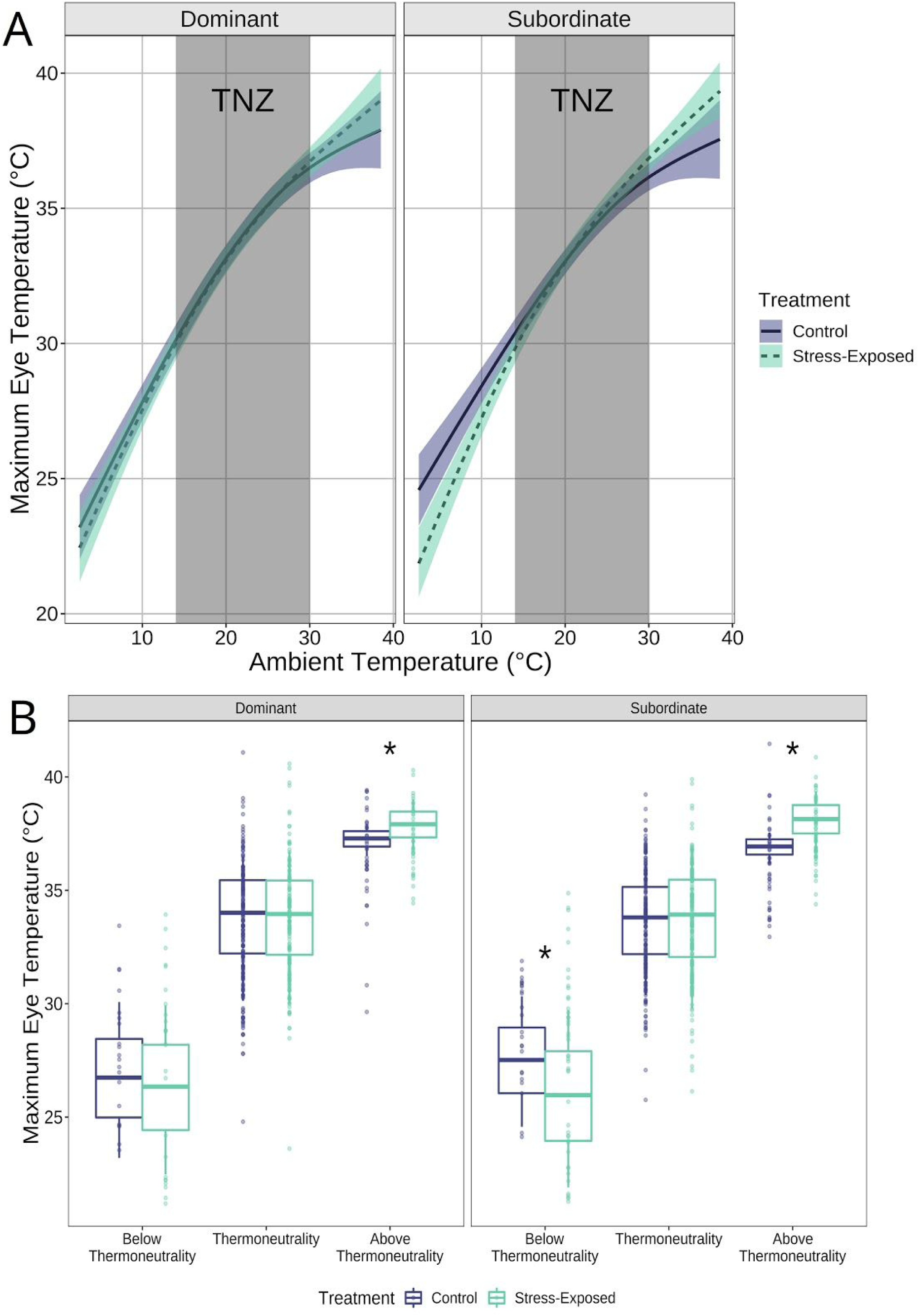
Effects of repeated stress exposure on eye region temperature (mean, hourly) of socially dominant and socially subordinate Black-capped Chickadees across ambient temperature. **A** | Eye temperature measurements (° *C*) calculated from infra-red thermographic images that were captured from 20 Black-capped Chickadees across sixty days (n_images_ = 1027, n_socially dominant_ = 8; n_socially subordinate_ = 12). Solid lines and dashed lines represent mean eye region temperature of individuals under control and stress exposure treatments, respectively (marginal means, calculated from a generalised additive mixed effects model). Stress exposure significantly influenced the relationship between ambient temperature and eye region temperature in social subordinates alone (*p* = 0.026). Purple and green bands represent 95% confidence intervals around mean estimates for control and stress exposure treatments, respectively. Grey rectangles represent the estimate thermoneutral zone (“TNZ”) of Black-capped Chickadees (14° *C* - 30° *C*). **B** | Distribution of eye region temperature measurements (° *C*) **from socially dominant and socially subordinate Chickadees at** ambient temperatures below, at, and above estimated thermoneutrality. Lower and upper ranges of boxes represent the 1_st_ and 3_rd_ quartiles respectively, whilst whiskers represent ± 1.58 *x* the interquartile range. Purple and green dots represent raw eye region temperature measurements at control and stress exposure treatments respectively. Asterisks represent significant differences between treatments at an ambient temperature zone (*p <* 0.05).

Interestingly, neither treatment nor social status independently explained surface temperature in our study animals (mean surface temperature = 33.4°C regardless of categorical grouping; treatment: *p* = 0.706; social status: *p* = 0.795; Tab. 1); an interaction between each factor was also insufficient to explain surface temperature profiles (*p* = 0.726; Tab. 1). The combined effects of time of day and flight enclosure orientation (i.e. east or west facing; n = 2 per cardinal direction) were, however, significantly correlated with surface temperature (time of day ⊗ flight enclosure: *p* <0.001; Tab. 1), and this diel rhythm differed according to treatment type (treatment : time of day ⊗ enclosure orientation: *p* <0.001; Tab. 1). A main effect of sex on surface temperature was not significant (mean surface temperature of males ± s.d. = 33.2°C ±3.45, mean surface temperature of females ± s.d. = 33.7°C ± 3.45; *p* = 0.895; Tab. 1).

### Social status influences the effects of stress exposure on dry heat transfer

Similar to our results regarding surface temperature, total dry heat transfer (q_Tot_) was significantly correlated with ambient temperature (*p* < 0.001; Tab. 2). This relationship between ambient temperature and q_Tot_ was significantly influenced by stress exposure in social subordinates alone (treatment ⊗ ambient temperature: *p* = 1.000; status : treatment ⊗ ambient temperature: *p* = 0.004; Tab. 2; Fig. 4), as predicted by the Thermoprotective Hypothesis. At ambient temperatures below thermoneutrality, q_Tot_ of social subordinates significantly fell during stress exposure treatments (mean Δ ± s.e.m. = -4.619 mW ± 1.450, *t*_*1005*_ = 3.190, *p* = 0.002); a similar fall was not observed among social dominants (mean Δ ± s.e.m. = −0.023 mW ± 1.220, *t*_*1005*_ = 0.019, *p* = 0.985). Interestingly, at ambient temperatures above thermoneutrality, stress exposure did not significantly influence q_Tot_ in socially dominant or socially subordinate individuals (social subordinates: mean Δ ± s.e.m. = 2.381 mW ± 1.440, *t*_*1005*_ = 1.650, *p* = 0.099; social dominants: mean Δ ± s.e.m = −0.03 mW ± 1.570, *t*_*1005*_ = 0.014, *p* = 0.989; Fig. 4), however, our analyses detected a trend of q_Tot_ increasing in response to stress exposure treatments among social subordinates alone (*p* < 0.1). Neither social dominants nor social subordinates displayed significant differences in q_Tot_ at ambient temperatures at thermoneutrality (social subordinates: mean Δ ± s.e.m. = 0.225 mW ± 0.536, *t*_*1005*_ = 0.420, *p* = 0.675; social dominants: mean Δ ± s.e.m. = −0.023 mW ± 0.584, *t*_*1005*_ = 0.039, *p* = 0.969; Fig. 4).

**Table 2.** **Results of a GAMM testing the influence of social status on total dry heat loss (qTot) across the eye region of Black-capped Chickadees (n = 20) across control and stress-exposed conditions**. q_Tot_ was measured across the eye region of Chickadees (n_dominant_ = 8; n_subordinate_ = 12) across 60 days (n_days_ = 30 per treatment type) and represent means per hour. Coefficient estimates (β for linear predictors, and mean β, 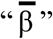 for smooth predictors) and standard errors around coefficient estimates (s.e.m.) are presented for each fixed-effect predictor. Degrees of freedom (*d*.*f*.) are reported for linear predictors, and estimated degrees of freedom (*e*.*d*.*f*.) are reported for smooth predictors. Asterisks (*) signify a significant effect at α = 0.05, and ⊗ represents a tensor product; n_measurements_ = 458; social subordinates: n_measurements_ = 569. This model explained 84.870% of deviance in surface temperature data.

| Linear Predictors |  |  |  |  |  |
| --- | --- | --- | --- | --- | --- |
| Predictor | Coefficient ($\beta$) | s.e.m. | *d.f.* | *t* | *p* |
| Intercept | 26.846 | 1.492 | 1 | 17.993 | <0.001* |
| Social Status | -0.127 | 0.488 | 1 | -0.259 | 0.796 |
| Sex | 0.091 | 0.737 | 1 | 0.124 | 0.901 |
| Stress Exposure | 0.230 | 0.912 | 1 | 0.252 | 0.801 |
| Stress Exposure:Status | 0.104 | 0.345 | 1 | 0.302 | 0.763 |
| Smooth Predictors |  |  |  |  |  |
| Predictor | Mean Coefficient ($\bar{\beta}$) | Mean s.e.m. | *e.d.f.* | *F* | *p* |
| Ambient Temperature | -19.600 | 1.310 | 2.261 | 103.745 | <0.001* |
| Ambient Temperature $\otimes$ Stress Exposure | 0.000 | 1.800 | 3.000 | <0.001 | 1.000 |
| Social Status: Ambient Temperature $\otimes$ Stress Exposure | -0.710 | 1.470 | 3.051 | 1.883 | 0.004* |
| Time of Day $\otimes$ Enclosure Orientation | -0.047 | 1.590 | 5.454 | 7.039 | <0.001* |
| Stress Exposure : Time of Day $\otimes$ Enclosure Orientation | -0.273 | 1.550 | 3.410 | 5.283 | <0.001* |
| Random Effects |  |  |  |  |  |
| Predictor | Variance |  |  |  |  |
| Bird Identity | 1.258 |  |  |  |  |
| Date of Photo | 2.223 |  |  |  |  |
| Enclosure Identity | 2.376 |  |  |  |  |

**Figure 4.**
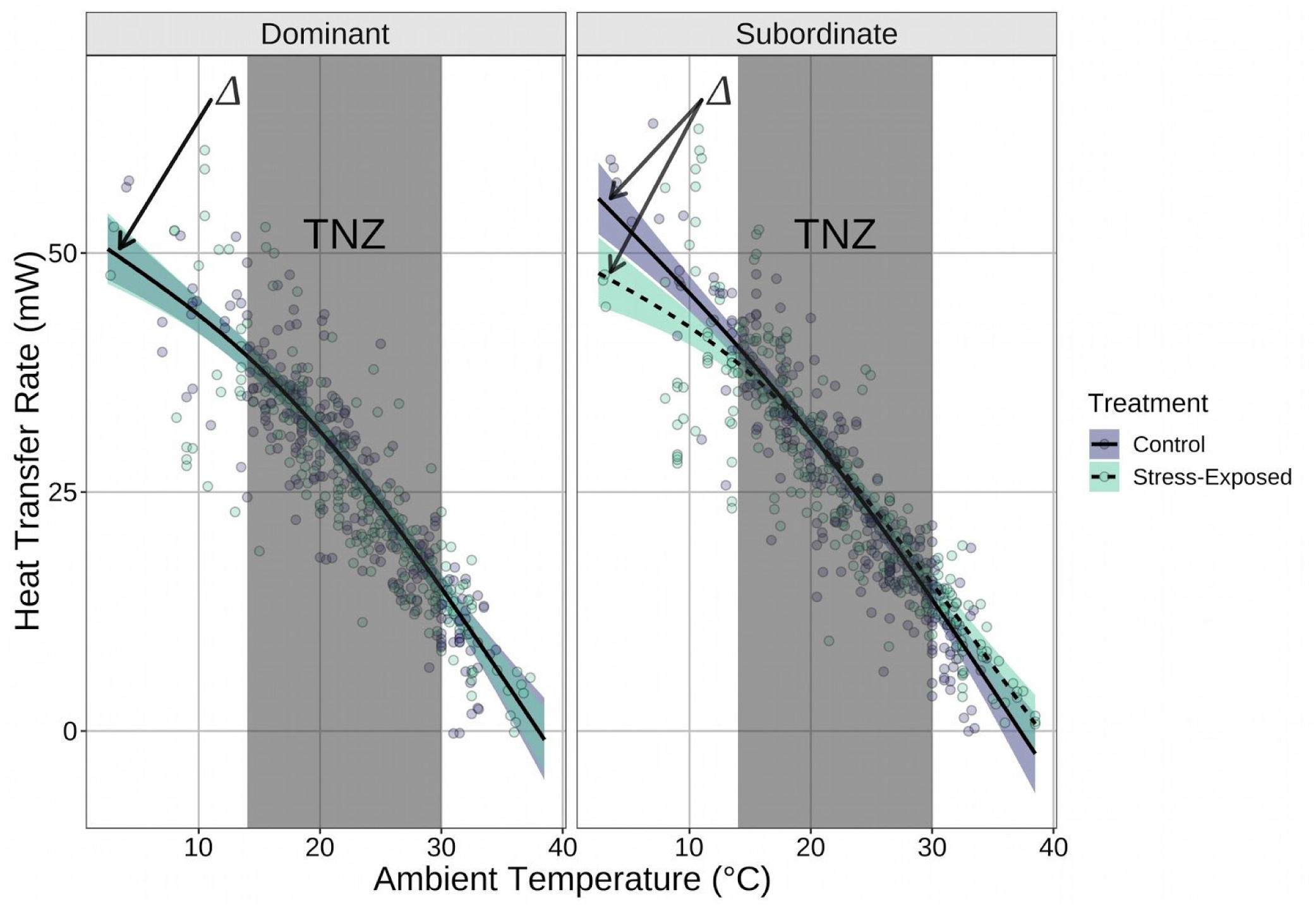
Dry heat transfer across ambient temperature in Black-capped Chickadees, as influenced by stress exposure and social status. Mean dry heat transfer (q_Tot_; marginal mean) across the eye region was calculated for socially dominant and socially subordinate individuals at naturally occurring ambient temperature, and during stress exposure and control treatments from a generalised additive model (n_socially dominant_ = 8, n_socially subordinate_ = 12, n_days_ = 60, n_observations_ = 1027). Dashed lines represent mean trends under control treatments and solid lines represent mean trends under stress exposure treatments; purple and green bands represent 95% confidence intervals around control treatment and stress exposure treatment trends respectively. Delta symbols denote curves between which comparisons were tested. Grey rectangles represent the estimated thermoneutral zone (“TNZ”) of Black-capped Chickadees (14° *C* - 30° *C*). Purple and green dots represent raw heat transfer measurements at control and stress exposure treatments respectively. Among social dominants, the relationship between ambient temperature and dry heat transfer did not significantly differ between control and stress exposure treatments (*p* = 0.112; α = 0.05). Contrastingly, social subordinates decreased dry heat loss at low ambient temperatures and increased dry heat loss at high ambient temperatures under stress exposure, when compared with controls (*p* < 0.001; α = 0.05).

We did not detect a significant main effect of treatment (mean q_Tot_ ± s.d.: control treatments = 25.600 mW 12.100; stress exposure treatments = 25.100 mW ± 12.200; *p* = 0.480; Tab. 2), social status (mean q_Tot_ ± s.d.: social subordinates = 25.200 mW 12.500; social dominants = 25.600 mW ± 11.800; *p* = 0.502; Tab. 2) or sex (mean q_Tot_ ± s.d.: males = 25.600 mW ± 11.40; females = 25.100 mW ± 13.000; *p* = 0.578; Tab. 2) on q_Tot_ from the eye region, paralleling our results with respect to surface temperature. Furthermore, an interactive, linear effect of social status and treatment on qTot (irrespective of ambient temperature) was not detected (*p* = 0.577; Tab. 2). Similar to our analysis of surface temperature, the combined effects of time of day and enclosure orientation significantly predicted q_Tot_ (time of day ⊗ flight enclosure; *p* <0.001; Tab. 2), and this relationship was significantly influenced by treatment type (treatment : time of day ⊗ flight enclosure: *p* <0.001; Tab. 2).

## 4. Discussion

### 4.1 Surface temperature responses to stress exposure represent an energetic trade-off

Stress-induced changes in surface temperature have previously been viewed as evolutionary “spandrels” that emerge as a by-product of experimental design, or vasomotor responses that serve alternative functional purposes (e.g. Andreasson et al., 2019; Jerem et al., 2015, 2018). Here, however, we provide evidence that thermal responses to stress exposure may emerge from a trade-off between thermoregulation and stress responsiveness, whereby individuals that are resource-limited (in our experiment, socially subordinate individuals; Supplemental Results; SFig. 1-2) seek to reduce their investment in thermoregulation when confronted with perceived stressors (the Thermoprotective Hypothesis). Indeed, our results show that during exposure to repeated stressors, social subordinates experienced a greater reduction in surface temperature at low ambient temperatures (e.g. those below thermoneutrality), and a greater increase in surface temperature at high ambient temperatures (e.g. those above thermoneutrality) than social dominants (Fig. 3; our first prediction) when compared with themselves under control conditions. Furthermore, this exacerbated surface temperature response to stressors appeared to endow social subordinates with a larger decrease in heat loss (a reduced q_Tot_), and larger increase in heat dissipation (an increased q_Tot_;, albeit, non-significant) during stress exposure than social dominants in cold and warm environments respectively (Fig. 4; our second prediction). At ambient temperatures below thermoneutrality, for example, social subordinates reduced their q_Tot_ across the eye region by an average of 4.62 mW (Results and SFig. 1; approximately 12% of maximal dry heat transfer; 54.4 mW) during stress exposure - a reduction in q_Tot_ that is substantially larger those experienced by social dominants at equivalent ambient temperatures (−0.02 mW; Results and SFig. 1). A similar disparity in q_Tot_ between social dominants and social subordinates was observed at ambient temperatures above thermoneutrality (social subordinates: Δ q_Tot_ ≈ 2.38 mW, social dominants: Δ q_Tot_ ≈ −0.03 mW).

For stress exposed social subordinates, our observed reduction in q_Tot_ by 4.62 mW in the cold, and increase in q_Tot_ by 2.38 mW in the heat under stress exposure corresponds with energetic savings of approximately 1.7%, and 0.9% of basal metabolic rate respectively, compared with rested social subordinates (as estimated for a 10.5 g individual, according to Petit, Lewden, & Vézina, 2013). Although such savings are arguably small, it is likely that stress-induced changes in surface temperature and q_Tot_ extend beyond the eye region alone, therefore rendering our perceived energetic savings underestimates of true, total savings experienced across the whole animal. In most birds, for example, the majority of environmental heat transfer occurs at exposed integument beyond the eye region (i.e. the legs and bill: Steen & Steen, 1965; Tattersall, Arnaout, & Symonds, 2017; Ward et al., 1999); thus, cumulative effects of stress exposure on q_Tot_ and subsequent energetic savings are therefore likely to exceed those experienced at the eye region alone, if thermal responses to stress exposure are consistent across exposed integument (as supported by stress-induced declines in wattle and comb temperature of Domestic Chickens, *Gallus gallus*, held at sub-thermoneutral temperatures, 18°C: Herborn et al., 2015; limits to thermoneutrality of Domestic Chickens, approximately 21°C: Barott & Pringle, 1946; Meltzer, 1983). Unfortunately, however, neither the legs nor bill were readily visible in our thermographic images, limiting our ability to explore stress-induced changes in heat transfer and energy allocation at the whole-animal level.

Interestingly, although social subordinates, but not social dominants, experienced a fall in mass across stress exposure treatments (*p* = 0.023; Supplemental Results; SFig. 2), the condition of individuals did not significantly differ according to social position upon termination of stress exposure treatments (residuals of mass regressed against wing-chord; *p* = 0.648; Supplemental Results; SFig. 3). In Siberian Hamsters (*Phodopus sungorus*), similar results have been reported whereby acute signals of negative energy balance (short-term glucoprivation) but not sustained energy deprivation (prolonged glucoprivation) modulated reproductive capacity of females (Carlton, Copper, & Demas, 2014). These results suggest that exacerbated thermal responses to stress exposure among social subordinates cannot simply be explained by comparative inanition in social subordinates, but rather, seem to reflect a more complex monitoring of energy balance and resource allocation in Chickadees. Additionally, whilst feeding rate was significantly elevated in social dominants when compared with subordinates (*p* = 0.005; Supplemental Results; SFig. 1), thermal consequences of hypophagia alone (i.e. a reduction in the time-averaged heat increment of feeding) are unlikely to explain differences in stress-induced thermal responses across social hierarchies. On the contrary, surface temperature of social subordinates did not differ from those of social dominants at thermoneutral temperatures, and even exceeded those of social dominants at supra-thermoneutral temperatures - neither of which would be predicted by an elevated heat increment of feeding among social dominants.

### 4.2 Magnitude but not presence of stress-induced thermoregulatory trade-offs depend upon thermal responses at the core

Among our experimental individuals, thermal responses to stress exposure were monitored at surface tissues alone. We therefore cannot elucidate whether differences in the magnitude of thermal responses between social dominants and social subordinates were driven by differences in peripheral vasomotion, or differences in core temperature fluctuations. Although neither possibility negates energetic benefits obtained by the responder (here, socially subordinate individuals), each likely hold differing consequences with respect to individual fitness, and total energetic savings attributed to stress-induced thermal responses.

At temperatures below thermoneutrality, declines in core body temperature can reduce the metabolic demand of Black-Capped Chickadees by as much as 10% (Reinertsen & Haftorn, 1986), but in doing so, enzymatic activity and neuromuscular response times are also likely to fall (discussed in Brodin, Nilsson, & Nord, 2017), owing to Q_10_ effects. In both Morning Dove (*Zenaida macroura*) and Blue Tits (*Cyanistes caeruleus*), such declines in neuromuscular response times during hypothermia have been anecdotally and empirically supported, with hypothermic individuals exhibiting reduced flight capacity when compared with normothermic conspecifics (Carr & Lima, 2013; Haftorn, 1972). Consequently, whilst permitting core body temperature to fall under stress exposure and resource limitation may be an energetically favourable strategy in the cold, long-term fitness consequences with respect to predator avoidance may be high. Contrastingly, although energetic savings acquired by reducing surface temperature under stress exposure are limited when compared with those obtained by reducing core temperature (1.7% vs. 10% of basal metabolic rate respectively; estimated according to Brodin et al., 2017; Petit et al., 2013), possible fitness consequences of doing so are nearly absent at ecologically relevant temperatures.

Similar to stress-induced thermal response in cold, those observed among social subordinates at high ambient temperatures (i.e. supra-thermoneutral; Fig. 3) are also likely to provide energetic advantages irrespective of whether they originate at core tissues, or are isolated to the periphery. At the level of core tissues, for example, permitting hyperthermia at high ambient temperatures may allow an individual that is resource-limited to both economise on available water, and minimise energetic expenses that accompany the act of cooling itself, both evaporatively and behaviourally. Such “adaptive hyperthermia” is a widely used strategy among desert bird species that contend with chronic resource limitations (reviewed in McKechnie & Wolf, 2019), and may well have been employed by social subordinates in our experiment when exposed to rotating stressors. Akin to when hypothermia is employed for resource conservation (discussed above), however, employing hyperthermia at the core for resource conservation is also likely to bear long-term fitness consequences in endotherms. In poultry, for example, heat stress has been shown to increase the production of reactive oxygen species (Azad, Kikusato, Hoque, & Toyomizu, 2010), which in turn, may have direct consequences on longevity (Monaghan, Metcalfe, & Torres, 2009). In contrast, although energetic savings obtained by increasing surface temperature at high ambient temperature may be low when compared with those obtained by concurrently elevating core tissue temperatures, long-term fitness consequences of doing so are probably negligible.

### 4.3 Physiology of social dominance alone is unlikely to explain differences in stress-induced thermal responses

A reduction in access to resources among social subordinates appears to be the most parsimonious explanation for their exacerbated surface temperature responses to stress exposure. However, it is possible that social subordination *per se* may influence the degree to which physiological processes underpinning the stress-induced thermal response occur. Stress-induced changes in body temperature occur rapidly, and have long been thought to be driven by activation of the sympathetic nervous system (discussed in Jerem et al., 2018). Studies addressing the influence of social status on indirect markers of sympathetic responsiveness (e.g. activation of the hypothalamic-pituitary-adrenal axis; reviewed in Sapolsky et al., 2000), however, provide little support for this explanation. In Mountain Chickadees (*Poecile gambeli*), for example, socially subordinate individuals display lower maximal corticosterone release in response to stress exposure than socially dominant individuals (Pravosudov, Mendoza, & Clayton, 2003) - a trend that has since been mirrored in Mallards (*Anas platyrhynchos*) and Northern Pintails (*Anas acuta*; Poisbleau, Fritz, Guillon, & Chastel, 2005). Broadly, sympathetic activation is thought to facilitate corticosterone release and activity (reviewed in Sapolsky et al., 2000), suggesting that socially subordinate individuals should therefore display reduced sympathetic function in addition to reduced corticosterone secretion. Such trends directly contradict our observations of increased stress-induced surface temperature responses among social subordinates. This suggests that generalised differences in sympathetic physiology correlated with social status are unlikely to explain our results.

## 5. Conclusion

In this study, we provide evidence that stress-induced changes in surface temperature emerge from a trade-off between stress responsiveness and thermoregulation. Specifically, we show that when access to resources is constrained by natural processes (here, social competition), individuals exposed to repeated stressors decrease their dry heat loss at low ambient temperatures, and increase their dry heat loss at high ambient temperatures (albeit non-significantly) when compared with controls. Such changes in heat transfer are likely to provide benefits by reducing demand on thermogenesis in the cold, or evaporative cooling in the heat. An understanding of the precise degree to which thermoregulatory expenses are offset under stress exposure and resource constraint, however, remains unclear, and will require a more holistic knowledge of regional body temperature changes. Together, our results provide further support for a functional value of stress-induced changes in surface temperature that, to date, has been largely unexplored. More importantly, however, our findings raise concerns about the capacity of endotherms to cope with the combined challenges of rising climatic instability and frequency of anthropogenic disturbances.

## Data Availability Statement

All data and R code used for the construction of this study are available at: https://datadryad.org/stash/share/QzmRV875_S5IYpMT0jxOYIsoO-ua0T3LX6AA9u-XpdQ.

## Competing Interests

No competing interests with respect to this research are declared.

## Funding

All funding for this research was provided by the Natural Sciences and Engineering Research Council (NSERC) of Canada (Grant #: RGPIN-04158-2014), and by an NSERC Collaborative Research and Training Experience Program (Grant #: CREATE 481954-2016).

## Figures

**Supplemental Figure 1.**
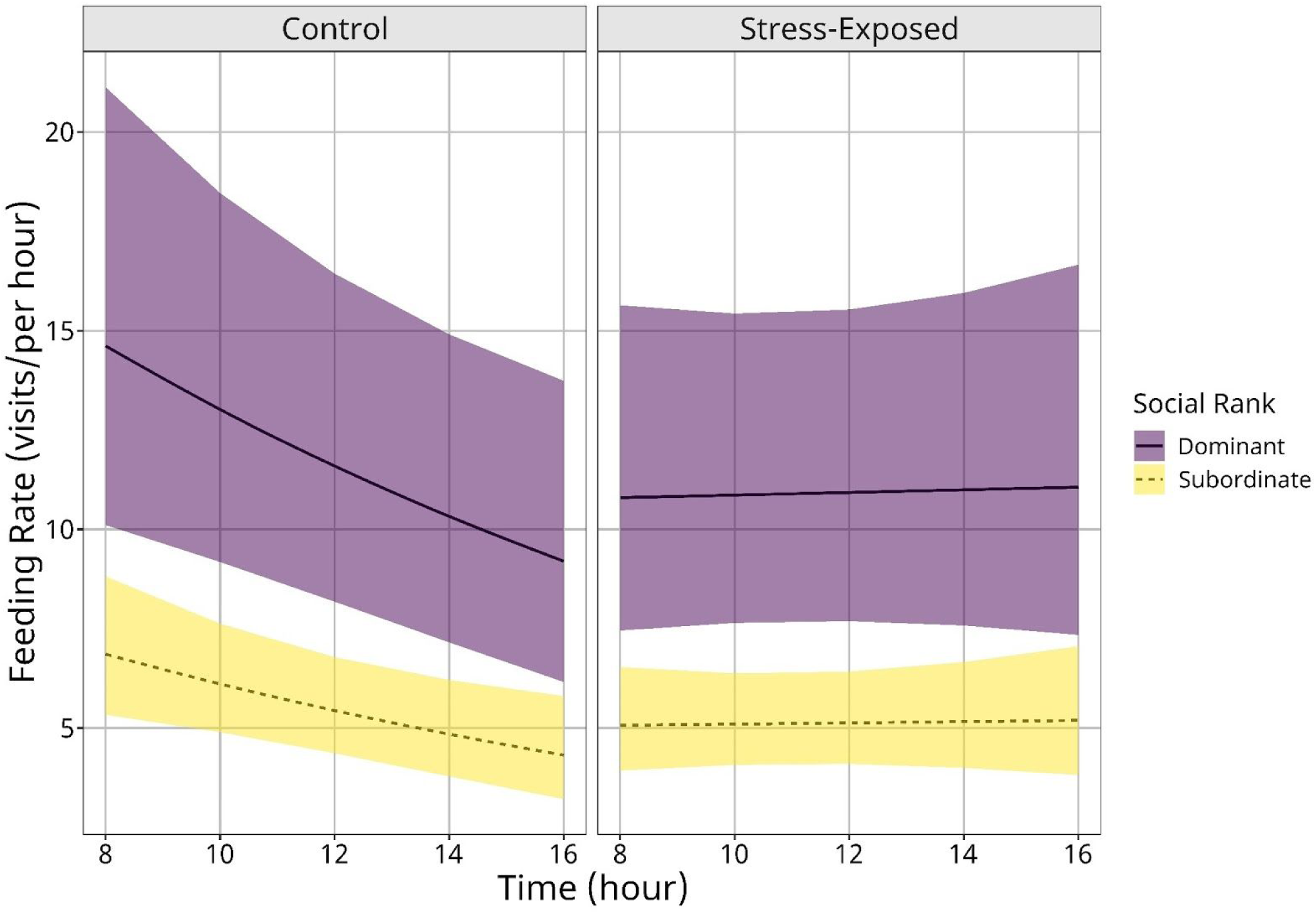
Effects of social status and stress exposure on feeding rate of Black-capped Chickadees across time of day. Feeding rate (visits/hour; marginal mean from generalised additive mixed effects model) was calculated from video observations made across 60 days, with a minimum of 1 hour of observation per flight enclosure (n_flight enclosures_ = 4), where individuals were identified by coloured leg bands. Solid lines represent mean feeding rate of socially dominant individuals and dashed lines represent that of socially subordinate individuals. Purple and yellow bands represent 95% confidence intervals around feeding rate estimates for socially dominant and socially subordinate individuals respectively. Feeding rate significantly differed between treatments (*p* = 0.007), and across social hierarchies (*p* = 0.001), at α = 0.05.

**Supplemental Figure 2.**
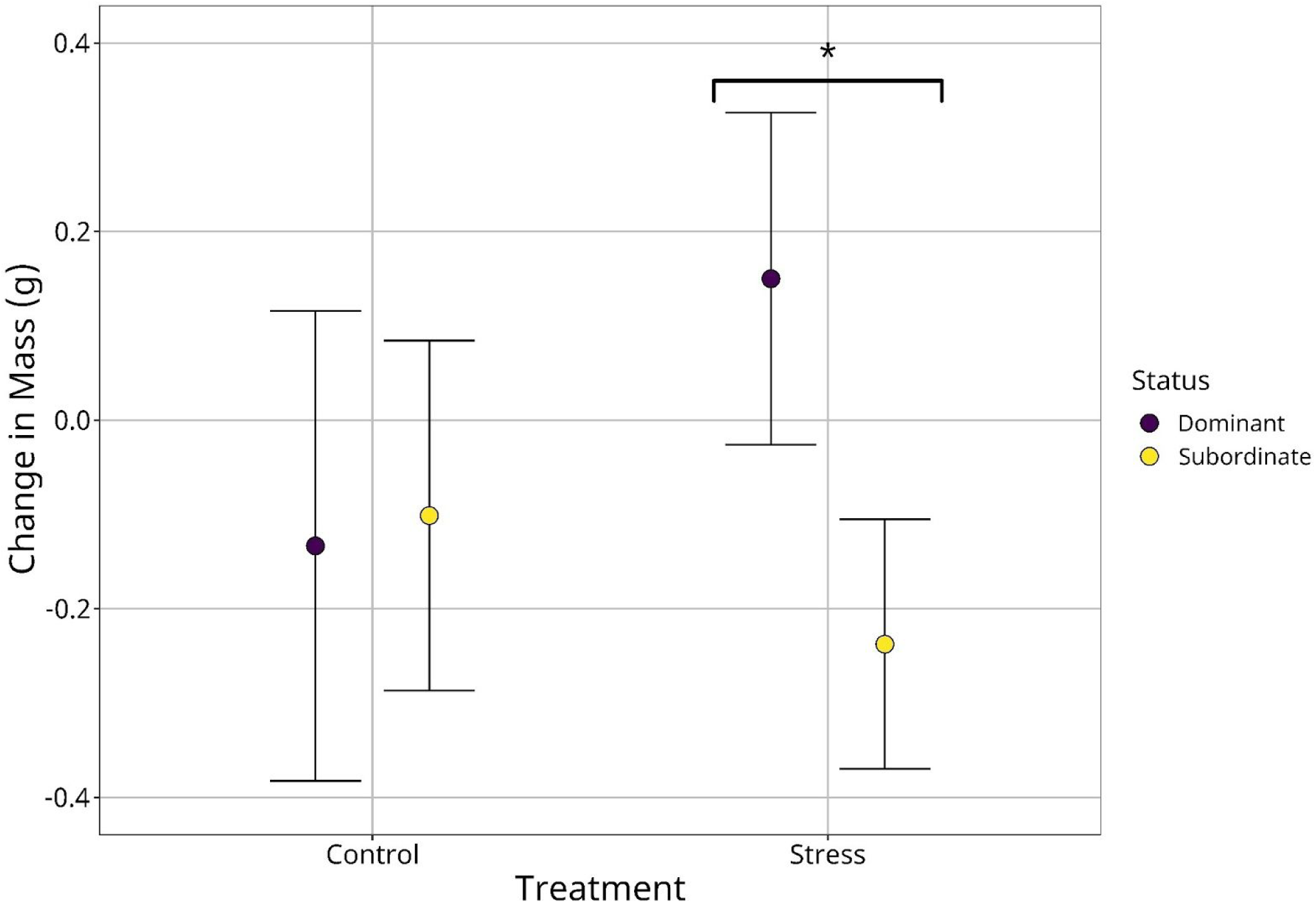
Effect of experimental treatment type on change in mass (across 30 days) of socially dominant and socially subordinate individuals. Mass (g) of each individual (n_socially dominant_ = 6; n_socially subordinate_ = 11) was measured at the onset and completion of each treatment. Purple dots and yellow dots represent mean change in mass (marginal means from linear mixed effects model) of socially dominant and socially subordinate individuals respectively. Socially subordinate individuals lost significantly more mass that socially dominant individuals across stress exposure treatments according to planned comparison (indicated by an asterisk; *p* = 0.023; α = 0.05). Whiskers represent 95% confidence intervals around means.

**Supplemental Figure 3.**
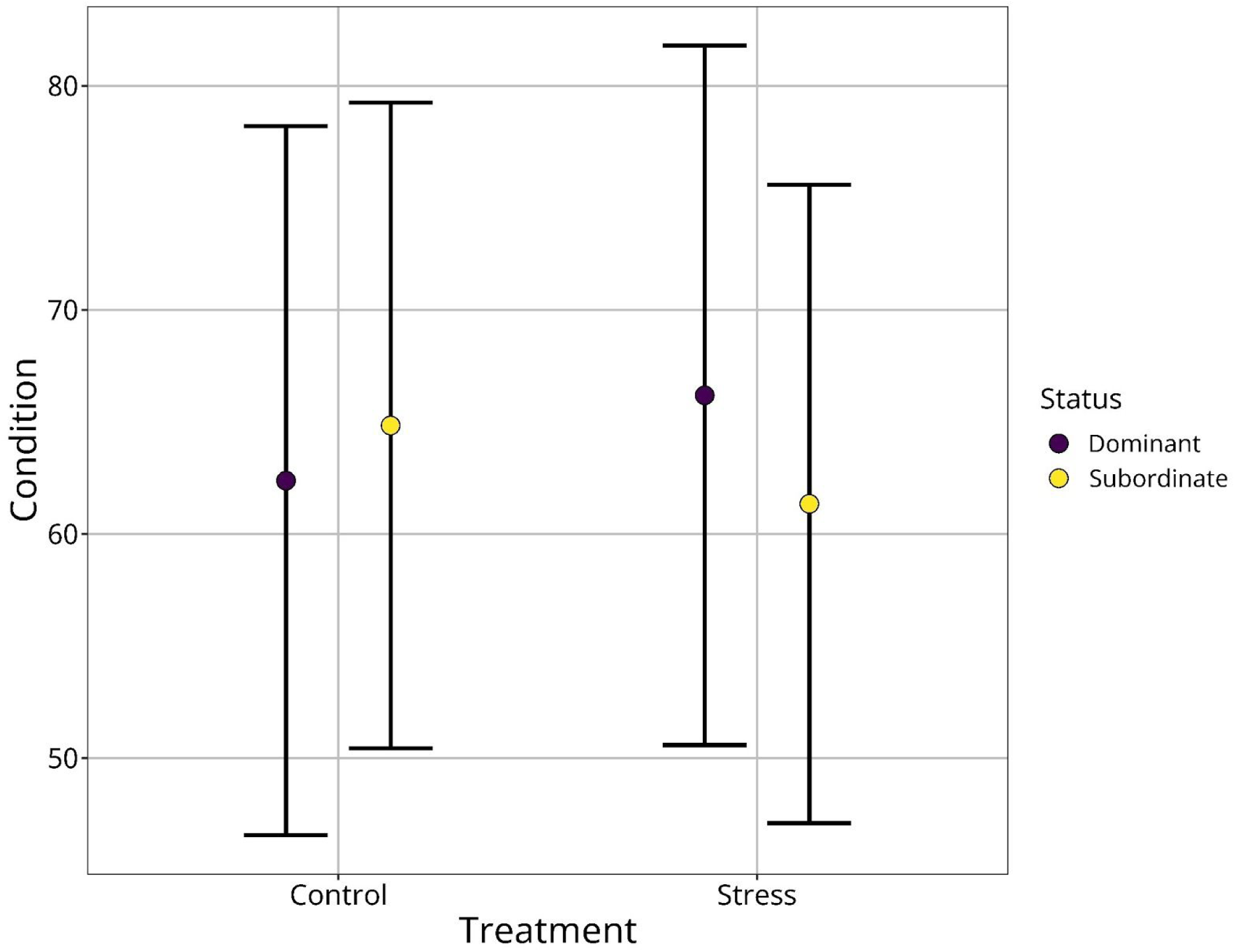
Effect of experimental treatment type on condition of socially dominant and socially subordinate individuals. Condition of each individual (n_socially dominant_ = 6; n_socially subordinate_ = 11) was calculated as the normalised residuals (0-100) from a linear model regression mass (g) against wing chord (mm). Purple dots and yellow dots represent mean condition (marginal means from linear mixed effects model) of socially dominant and socially subordinate individuals respectively, and whiskers represent 95% confidence intervals around means. No significant difference was detected between the condition of socially dominant and socially subordinate individuals following control or stress exposure conditions according to a linear mixed effects model and planned comparison (Control: *p* = 0.817; stress exposure: *p* = 1.000).

## References

Abolins-Abols, M., Hope, S., & Ketterson, E. (2016). Effect of acute stressor on reproductive behavior differs between urban and rural birds. Ecol Evol, 6(18), 6546–6555.

Aghajani, M., Vaez Mahdavi, M., Khalili Najafabadi, M., Ghazanfari, T., Azimi, A., & S Mahdi Dust, S. Arbab Soleymaniand. (2013). Effects of dominant/subordinate social status on formalin-induced pain and changes in serum proinflammatory cytokine concentrations in mice. PLoS One, 8(11), e80650.

Andreasson, F., Nord, A., & Nilsson, J. (2019). Body temperature responses of great tits *Parus major* to handling in the cold. Ibis.

Azad, A., Kikusato, M., Hoque, A., & Toyomizu, M. (2010). Effect of chronic heat stress on performance and oxidative damage in different strains of chickens. J Poult Sci, 47(1), 333–337.

Barott, H., & Pringle, E. (1946). Energy and gaseous metabolism of the chicken from hatch to maturity as affected by temperature: Four figures. Nutr J, 31(1), 35–50.

Bartlett, W. (1912). An experimental study of the arteries in shock. J Exp Med, 15(4), 415–428.

Best, R., & Fowler, R. (1981). Infrared emissivity and radiant surface temperatures of canada and snow geese. J Wildl Manag, 45(4), 1026–1029.

Breuner, C.W., & Berk, S.A. (2019). Using the van Noordwijk and de Jong resource framework to evaluate glucocorticoid-fitness hypotheses. Int Comp Biol, 59(2), 243–250.

Brodin, A., Nilsson, J., & Nord, A. (2017). Adaptive temperature regulation in the little bird in winter: Predictions from a stochastic dynamic programming model. Oecologia, 185(1), 43–54.

Burness, G., Armstrong, C., Fee, T., & Tilman-Schnidel, E. (2010). Is there an energetic-based trade-off between thermoregulation and the acute phaseresponse in zebra finches? J Exp Biol, 213(1), 1386–1394.

Cabanac, M., & Gosselin, F. (1993). Emotional fever in the lizard *callopistes maculatus* (teiidae). Anim Behav, 46(1), 200–200.

Calisi, R., Rizzo, N., & Bentley, G. (2008). Seasonal differences in hypothalamic egr-1 and gnih expression following capture-handling stress in house sparrows (*Passer domesticus*). Gen Comp Endocrinol, 157(3), 283–287.

Cannon, W. (1932). The wisdom of the body. New York, USA: Norton; Company Inc.

Carlton, E., Copper, C., & Demas, G. (2014). Metabolic stressors and signals differentially affect energy allocation between reproduction and immune function. Gen Comp Endocrinol, 208(1), 21–29.

Carr, J., & Lima, S. (2013). Nocturnal hypothermia impairs flight ability in birds: A cost of being cool. P Roy Soc B, 280(1772), 20131846.

Dymond, K., & Fewell, J. (1998). Gender influences the core temperature response to a simulated open field in adult guinea pigs. Physiol Behav, 65(4-5), 889–892.

Elo, A. (1978). The rating of chessplayers, past and present. New York, USA: Arco Pub.

Evans, J., Devost, I., Jones, T., & Morand-Ferron, J. (2018). Inferring dominance interactions from automatically recorded temporal data. Ethol, 124(3), 188–195.

Ficken, M., Weise, C., & Popp, J. (1990). Dominance rank and resource access in winter flocks of black-capped chickadees. Wilson Bull, 102(1), 623–633.

Fridolfsson, A., & Ellegren, H. (1999). A simple and universal method for molecular sexing of non-ratite birds. J Avian Biol, 30, 116–121.

Glase, J. (1973). Ecology of social organization in the black-capped chickadee. Living Bird, 12(1), 235–267.

Grachev, P., Li, X., & O’Byrne, K. (2013). Stress regulation of kisspeptin in the modulation of reproductive function. In A. Kauffman & J. Smith (Eds.), Kisspeptin signaling in reproductive biology (pp. 431–454). New York, USA: Springer.

Greenacre, C., & Lusby, A. (2004). Physiologic responses of amazon parrots (amazona species) to manual restraint. J Avian Med Surg, 18(1), 19–23.

Griffiths, R., Dean, S., & Dijkstra, C. (1996). Sex identification in birds using two chd genes. Proc R Soc Lond B, 263(1374), 1251–1256.

Grossman, A., & West, G. (1977). Metabolic rate and temperature regulation of winter acclimatized black-capped chickadees *parus atricapillus* of interior alaska. Ornis Scand, 8(2), 127–138.

Haftorn, S. (1972). Hypothermia of tits in the arctic winter. Ornis Scand, 3(2), 153–166.

Herborn, K., Graves, J., Jerem, P., Evans, N., Nager, R., McCafferty, D., & McKeegan, D. (2015). Skin temperature reveals the intensity of acute stress. Physiol Behav, 152(1), 225–230.

Herborn, K., Jerem, P., Nager, R., McKeegan, D., & McCafferty, D. (2018). Surface temperature elevated by chronic and intermittent stress. Physiol Behav, 191(1), 47–55.

Jerem, P., Herborn, K., McCaferty, D., McKeegan, D., & Nager, R. (2015). Thermal imaging to study stress non-invasively in unrestrained birds. J Vis Exp, 1–10.

Jerem, P., Jenni-Eiermann, S., Herborn, K., McKeegan, D., McCaferty, D., & Nager, R. (2018). Eye region surface temperature reflects both energy reserves and circulating glucocorticoids in a wild bird. Sci Rep, 8(1), 1907.

Jerem, P., Jenni-Eiermann, S., McKeegan, D., McCafferty, D., & Nager, R. (2019). Eye region surface temperature dynamics during acute stress relate to baseline glucocorticoids independently of environmental conditions. Physiol Behav, 210(1), 112627.

Kataoka, N., Hioki, H., Kaneko, T., & Nakamura, K. (2014). Psychological stress activates a dorsomedial hypothalamus-medullary raphe circuit driving brown adipose tissue thermogenesis and hyperthermia. Cell Metab, 20(2), 346–358.

King, M., & Swanson, D. (2013). Activation of the immune system incurs energetic costs but has no effect on the thermogenic performance of house sparrows during acute cold challenge. J Exp Biol, 216(1), 2097–2102.

Kinsey-Jones, J., Li, X., Knox, A., Wilkinson, E., Zhu, X., Chaudhary, A., … O’Byrne, K. (2009). Down-regulation of hypothalamic kisspeptin and its receptor, kiss1r, mRNA expression is associated with stress-induced suppression of luteinising hormone secretion in the female rat. J Neuroendocrinol, 21(1), 20–29.

Kirby, E., Geraghty, A., Ubuka, T., Bentley, G., & Kaufer, D. (2008). Stress increases putative gonadotropin inhibitory hormone and decreases luteinizing hormone in male rats. Proc Nat Acad Sci, 106(27), 11324–11329.

Lenth, R., Singmann, H., & Love, J. (2018). Emmeans: Estimated marginal means, aka least-squares means. R Package Version, 1(1).

Martin, L., Scheuerlein, A., & Wikelski, M. (2003). Immune activity elevates energy expenditure of house sparrows: A link between direct and indirect costs? Proc R Soc B, 270(1), 153–158.

McCafferty, D., Gilbert, C., Paterson, W., Pomeroy, P., Thompson, D., Currie, J., & Ancel, A. (2011). Estimating metabolic heat-loss in birds and mammals by combining infrared thermography with biophysical modelling. Comp Biochem Physiol A Mol Integr Physiol, 158(3), 337–345.

McKechnie, A., & Wolf, B. (2019). The physiology of heat tolerance in small endotherms. Physiol, 34(5), 302–313.

McKechnie, A., Gerson, A., McWhorter, T., Smith, E., Talbot, W., & Wolf, B. (2016). Avian thermoregulation in the heat: Evaporative cooling in five australian passerines reveals within-order biogeographic variation in heat tolerance. J Exp Biol, 220(13), 2436–2444.

Meltzer, A. (1983). Thermoneutral zone and resting metabolic rate of broilers. Br Poult Sci, 24(4), 471–476.

Merlo, J., Cutrera, A., Luna, F., & Zenuto, R. (2014). PHA-induced inflammation is not energetically costly in the subterranean rodent *Ctenomys talarum* (tuco-tucos). Comp Biochem Physiol A Mol Int Physiol, 175(1), 90–95.

Minkina, W., & Dudzik, S. (2009). Infrared thermography errors and uncertainties. Chichester, UK: Wiley Press.

Monaghan, P., Metcalfe, N., & Torres, R. (2009). Oxidative stress as a mediator of life history trade-offs: Mechanisms, measurements and interpretation. Ecol Lett, 12(1), 75–92.

Nord, A., & Folkow, L. (2019). Ambient temperature effects on stress-induced hyperthermiain svalbard ptarmigan. Biol Open, 8(6), bio043497.

Nord, A., & Nilsson, J. (2019). Heat dissipation rate constrains reproductive investment in a wild bird. Funct Ecol, 33(2), 250–259.

Nord, A., Sandell, M., & Nilsson, J. (2010). Female zebra finches compromise clutch temperature inenergetically demanding incubation conditions. Funct Ecol, 24(5), 1031–1036.

Oka, T., Oka, K., & Hori, T. (2001). Mechanisms and mediators of psychological stress-induced rise in core temperature. Psychosom Med, 63(1), 476–486.

Ots, I., Kerimov, A., Ivankina, E., Ilyina, T., & Hőrak, P. (2001). Immune challenge affects basal metabolic activity in wintering great tits. Proc R Soc B, 268(1), 1175–1181.

Parr, L., & Hopkins, W. (2000). Brain temperature asymmetries and emotional perception in chimpanzees, Pantroglodytes. Physiol Behav, 71(3-4), 363–371.

Petit, M., Lewden, A., & Vézina, F. (2013). Intra-seasonal flexibility in avian metabolic performance highlights the uncoupling of basal metabolic rate and thermogenic capacity. PLoS One, 8(6), e68292.

Poisbleau, M., Fritz, H., Guillon, N., & Chastel, O. (2005). Linear social dominance hierarchy and corticosterone responses in male mallards and pintails. Horm Behav, 47(4), 485–492.

Pravosudov, V., Mendoza, S., & Clayton, N. (2003). The relationship between dominance, corticosterone, memory, and food caching in mountain chickadees (*Poecile gambeli*). Horm Behav, 44(2), 93–102.

Proctor, C., Freeman, E., & Brown, J. (2010). Influence of dominance status on adrenal activity and ovarian cyclicity status in captive african elephants. Zoo Biol, 29(2), 168–178.

R Core Team. (2019). R: A language and environment for statistical computing. r foundation for statistical computing, vienna, austria. https://www.R-Project.org/.

Reinertsen, R., & Haftorn, S. (1986). Different metabolic strate-gies of northern birds for nocturnal survival. J Comp Physiol B, 156(1), 655–663.

Rey, S., Huntingford, F., Boltana, S., Vargas, R., Knowles, T., & Mackenzie, S. (2015). Fish can show emotional fever: Stress-induced hyperthermia in zebrafish. Proc R Soc B, 282(1819), 20152266.

Rich, E., & Romero, L. (2005). Exposure to chronic stress downregulates corticosterone responses to acute stressors. Am J Physiol Regul Integr Comp Physiol, 288, R1628–R1636.

Rising, J., & Hudson, J. (1974). Seasonal variation in the metabolism and thyroid activity of the black-capped chickadee (*parus atricapillus*). Condor, 76, 198–203.

Robertson, J., Mastromonaco, G., & Burness, G. (2020). Evidence that stress-induced changes in surface temperature serve a thermoregulatory function. J Exp Biol, 223(4), jeb213421.

Romero, L., Dickens, M., & Cyr, N. (2009). The reactive scope model—a new model integrating homeostasis, allostasis, and stress. Horm Behav, 55(3), 375–389.

Sánchez-Tójar, A., Schroeder, J., & Farine, D. (2018). A practical guide for inferring reliable dominance hierarchies and estimating their uncertainty. J Anim Ecol, 87(3), 594–608.

Sapolsky, R. (2004). Why zebras don’t get ulcers: The acclaimed guide to stress, stress-related diseases, and coping-now revised and updated. New York, USA: Holt Publishing.

Sapolsky, R., Romero, L., & Munck, A. (2000). How do glucocorticoids influence stress responses? Integrating permissive, suppressive, stimulatory, and preparative actions. Endocrin Rev, 21(1), 55–89.

Schubert, K., Mennil, D., Ramsay, S., Otter, K., Boag, P., & Ratcliffe, L. (2007). Variation in social rank acquisition influences lifetime reproductive success in black-capped chickadees. Biol J Linnean Soc, 90(1), 85–95.

Selye, H. (1950). Stress and the general adaptation syndrome. BMJ, 1(4667), 1383.

Seutin, G., White, B., & Boag, P. (1991). Preservation of avian blood and tissue samples for dna analyses. Can J Zool, 9(1), 82–90.

Simpson, G. (2018). Modelling palaeoecological time series using generalised additive models. Front Ecol Evol, 6(1), 149.

Smith, S. (1976). Ecological aspects of dominance hierarchies in black-capped chickadees. Auk, 93(1), 95–107.

Smith, S. (1991). The black-capped chickadee. behavioral ecology and natural history. Ithica, USA: Cornell University Press.

Speakman, J. (2008). The physiological costs of reproduction in small mammals. Phil Trans R Soc B, 363(1), 375–398.

Steen, I., & Steen, J. (1965). The importance of the legs in the thermoregulation of birds. Acta Physiol Scand, 63(3), 285–291.

Stier, K., Almasi, B., Gasparini, J., Piault, R., Roulin, A., & Jenni, L. (2009). Effects of corticosterone on innate and humoral immune functions and oxidative stress in barn owl nestlings. J Exp Biol, 212(13), 2085–2091.

Svensson, E., R, L., Koch, C., & Hasselquist, D. (1998). Energetic stress, immunosuppression and the costs of an antibody response. Funct Ecol, 12(6), 912–919.

Tattersall, G. (2016). Infrared thermography: A non-invasive window into thermal physiology. Comp Biochem Physiol A Mol Integr Physiol, 202, 78–98.

Tattersall, G., Arnaout, B., & Symonds, M. (2017). The evolution of the avian bill as a thermoregulatory organ. Biol Rev, 92(3), 1630–1656.

van Oort, H., Otter, K., Fort, K., & McDonnell, Z. (2007). Habitat, dominance, and the phenotypic quality of male black-capped chickadees. Condor, 109(1), 88–96.

Vleck, C. (1981). Energetic cost of incubation in the zebra finch. Condor, 83(1), 229–237.

Ward, S., Rayner, J., Möller, U., Jackson, D., Nachtigall, W., & Speakman, J. (1999). Heat transfer from starlings *sturnus vulgaris* during flight. J Exp Biol, 202(12), 1589–1602.

Wood, S. (2011). Fast stable restricted maximum likelihood and marginal likelihood estimation of semiparametric generalized linear models. J R Stat Soc B, 73(1), 3–36.

Yeo, I. (2005). Hippocrates in the context of galen: Galen’s commentary on the classification of fevers in epidemics vi. Stud Anc Med, 31, 433–443.

Yokoi, Y. (1966). Effect of ambient temperature upon emotional hyperthermia and hypothermia in rabbits. J Appl Physiol, 21(6), 1795–1798.

