## Supplemental Material for "A temperate endotherm trades thermoregulation for self-preservation: stress-induced changes in surface temperature as thermoregulatory trade-offs"

### Materials and Methods

#### Thermographic and filming protocols

All thermographic filming protocols used in this study are described in Robertson, Mastromonaco, & Burness (2020). Briefly, because time of day is likely to influence body temperature in Chickadees (Reinertsen & Haftorn, 1984), we rotated thermographic filming among flight enclosures (cardinally clockwise) within each day, and initiated filming at a randomly determined flight enclosure each morning to control for the effect of time alone on surface temperature readings. Specifically, we captured thermographic images for approximately one hour at each flight enclosure, and following each filming interval, we rotated our thermographic camera to a new flight enclosure, then initiated a new filming interval.

In parallel to thermographic imaging, we digitally filmed individuals at feeding platforms to determine the identity of those present in thermographic images, and to monitor social interactions that occurred between individuals during feeding. Digital video was captured by placing a small digital camera (Action Cam™, Sony, Toronto, Ontario, CA) adjacent to our thermographic camera in a given camera-box, and fitting the lens to a perforation oriented toward a respective feeding platform. Digital filming schedules followed thermographic imaging schedules, such that our digital camera was rotated alongside our thermographic camera. In total, we collected 445 hours and 30 minutes of digital video (average time per flight enclosure = 111 hours and 23 minutes), across 60 days of filming. Upon completion of the experiment, digital video and thermographic images were aligned visually, and the identity of individuals present in thermographic images were determined by assessing coloured leg-band combinations in parallel, digital video frames.

#### Heat transfer calculations

In our calculations of convective heat transfer $q_{Conv}$ and radiative heat transfer $q_{Rad}$ from the eye region of Black-capped Chickadees, we followed methods described in (Robertson et al., 2020). Specifically, we allowed the thermal conductivity of air (W/m $^{\circ}$C$^{-1}$), the kinematic viscosity of air (m$^2$/S; at an assumed atmospheric pressure of 101.325 kPa), and the thermal expansion coefficient of air (1/K) to vary according to the ambient temperature experienced at the time of image capture. Kinematic viscosity and air conductivity calculations were conducted using the R packages "bigleaf" and "Thermimage" respectively (Knauer, El-Madany, Zaehle, & Migliavacca, 2018; Tattersall, 2019).

#### Social status determination

##### Enumeration of video observations

To test whether thermal responses to stress exposure differed between dominant and subordinate Black-capped Chickadees, we required estimation of linear dominance hierarchies within each experimental flight enclosure, and subsequently, individual social positions within each hierarchy. Typically, studies estimating dominance hierarchies from video observations rely upon subjective categorisations of behaviours as agonistic or non-agonistic (Silva et al., 2018; Snijders, Naguib, & van Oers, 2016) and consequently, are likely to suffer from observer bias. To eliminate observer bias from our estimates of dominance hierarchies, we chose to enumerate video observations as presence and absence data across time (per individual), then apply static and consistent criteria to infer and categorise social interactions without requiring visual observation of their occurrence (see "Social status determination"). Here, video observations were converted to enumerated time-line data by determining the presence of individuals at feeding platforms across video frames, then assigning each presence and absence per individual a binomial value (present = 1, absent = 0) for a given second of observation.

Presence of individuals at feeding platforms was determined by detecting arrivals and departures from video recordings using automated motion detection, then identifying individuals present using coloured leg-band combinations. Specifically, we scanned videos for the presence of motion (defined as a change in pixel content between adjacent frames exceeding 1.75%, at a frame rate of 8 fps) in R (R Core Team, 2019), then extracted still frames from $\pm 2$ seconds around the time of motion (8 fps; $n_{frames} = 16$). Individuals observed to be arriving to, present at, and departing from feeding platforms within extracted frames were then identified by observation of coloured leg band combinations, and arrival, presence, and departure times were documented according to real time (as determined from digital time-stamps of parallel thermographic images).

#### Categorization of behavioural interactions

In our study, we defined agonistic interactions as displacement and waiting events from feeding platforms (similar to Evans, Devost, Jones, & Morand-Ferron, 2018), whilst non-agonistic interactions included prolonged and simultaneous presence of multiple individuals at a single feeding platform. We defined displacement events as social interactions wherein the arrival of one individual to a feeding platform (individual A) stimulated the departure of another individual that was previously present at the same feeding platform (individual B), within one second of individual A's arrival. Research comparing social hierarchies that were estimated using radio frequency identification (RFID) and simultaneous video observations has shown that the correlation between rank estimates is highest when the criteria for displacement events are limited, such that the minimum length of time that individual A must remain at the feeding platform following its arrival ($j$) is 5 seconds (Evans et al., 2018). Because our feeding data were enumerated in a manner comparable to those obtained using RFID technology, identification of displacement events were limited such that $j$ must be equal to or greater than 5 seconds. We defined waiting events as social interactions wherein the arrival of individual A to a feeding platform was contingent upon the departure of individual B from the same feeding platform, and occurred between one and three seconds after the departure of individual B.

Unlike our criteria for displacement events, no minimum retention time of individual A was defined (i.e. $j > 0$ seconds). Non-agonistic encounters were defined as the simultaneous presence of two individuals at a feeding platform for more than 2 seconds.

To support calculation of a randomised Elo ratings across individuals, we assigned those present in agonistic interactions a win or loss according to whether they received priority access to feeding platforms; those who displaced other individuals or fed freely whilst others awaited their departure were assigned a win, and those subjected to displacement or waiting for access to feeding platforms were assigned a loss. Individuals present in non-agonistic interactions were assigned neutral "draws". To ensure that our Elo rating estimates were conservative, we assumed that displacement events held greater importance in defining an individual's rating than waiting events, given that feeding order and time may also be randomly determined. To do so, we weighted displacements events more heavily than waiting events in our calculation of Elo ratings by assigning each a sensitivity score ($K$) of 50, and 100 respectively.

Randomised Elo ratings were calculated for each individual by randomly sampling 20 days of observation data, then loading these data into the function "elo.seq" from the R package "EloRating" (Neumann & Kulik, 2019). This process was iterated 1000 times, and mean, final Elo ratings $\pm 95\%$ CIs were calculated per individual, per flight enclosure, across iterations. Chickadees within each flight enclosure were then ranked descending according to their mean Elo rating (rank 1 = most dominant, rank 5 = least dominant), then categorised as either socially dominant or socially subordinate. Here, two individuals with the highest mean Elo ratings within each enclosure were considered socially dominant ($n_{per\ enclosure}$ = 2; $n_{Total}$ = 8; Fig. 2), whilst the remaining individuals were considered socially subordinate ($n_{per\ enclosure}$ = 3; $n_{Total}$ = 12; Fig. 2). Partitioning of individuals in this way was chosen to balance sample size between status groups whilst minimising the number of individuals experiencing regular top-down antagonism in socially dominant categories.

#### Assessment of feeding rate and mass loss

Upon onset and completion of each experimentation treatment, individuals were weighed (g) to assess the influence of treatment on mass. Individuals displaying signs of distress (panting, eye closure), however, were excluded from mass measurements (n = 3 individuals; $n_{socially\ dominant}$ = 2, $n_{socially\ subordinate}$ = 1). To assess the influence of social status on feeding rate, we calculated the number of visits made to a feeding platform by each individual across each continuous hour of observation (n = 60 days of observation; derived from enumerated, presence/absence data; see "Social status determination: Enumeration of video observations").

#### Statistical Analyses

##### Effect of social status on feeding rate and stress-induced mass loss

In this study, we predicted that individuals that were resource-limited (socially subordinate individuals) would exhibit a greater reduction in surface temperature and dry heat-loss at low ambient temperatures, and a greater increase in surface temperature and dry heat loss at high ambient temperatures following stress exposure than that that were not resource limited (socially dominant individuals). Because this prediction is contingent upon differences in resource access being detectable across social hierarchies in our experiment, however, we further predicted that social dominance would have a positive influence on access to supplemental food in our study population. Additionally, we predicted that resource limitations (as measured by a decline in body mass) would be detectable in socially subordinate individuals, but not socially dominant individuals, following prolonged stress exposure.

To statistically test whether socially dominant individuals had greater access to supplied food than socially subordinate individuals, we used a generalised additive mixed effects model in the R package "mgvc" (Wood, 2011) with feeding rate as the response variable and social status as a binomial, parametric predictor. Previous studies have highlighted a non-linear effect of ambient temperature and time of day on the frequency of visits to supplemental food in Black-capped Chickadees (Bonter, Zuckerberg, Sedgwick, &

Hochachka, 2013; Brittingham & Temple, 1992). Consequently, we initially included both time of day (hour) and average ambient temperature per hour (°C) in our model as independent cubic regression splines, each with 4 knots to permit prediction of curvilinear relationships whilst circumventing model over-fit. Our results, however, provided little evidence for a curvilinear relationship between time of day and feeding rate within our window of observations, so our cubic regression spline modelling the relationship between time of day and feeding rate was removed, and time of day was subsequently included as a parametric, linear predictor. Because exposure to psychogenic stressors is capable of influencing feeding behaviour in birds (Favreau-Peigné et al., 2014), treatment (stress-exposed or control) was included as parametric predictor (binomial), and interactions between both treatment and time of day, and treatment and mean ambient temperature were also included to account for possible time and temperature dependant feeding patterns following stress exposure. Additionally, an interaction between social status and treatment was included to account for possible differences in behavioural responses to stress across hierarchies. To control for statistical non-independence between feeding rate measurements that were derived across the same day and/or same individual, both date (Julian) and bird identity were included as random intercepts. Similarly, capture locale (1 of 6) was also included as a random intercept to account for potential relatedness among individuals from the same place of capture, and possible population-specific physiological effects. Finally, we included sex as a binomial predictor to control for possible differences in feeding behaviour between male and female Chickadees. In our data, variance in feeding rate exceeded mean feeding rate ( $\Lambda > \mu$ ). We therefore assumed a negative binomial distribution with an estimated $\theta$ of 2.4 to capture over-dispersion.

#### Effect of treatment on individual mass, with respect to social status

We were interested in testing whether socially subordinate individuals, but not socially dominant individuals, were unable to maintain a constant body mass across stress exposure treatments. To do so, we used a linear mixed effects model ("LMM") in the R package "glmmTMB" (Brooks et al., 2017), with change in mass between the termination and onset of each treatment ($\Delta$g) as the Gaussian distributed response variable ($n_{individuals}$ = 17; $n_{socially\ dominant}$ = 6; $n_{socially\ subordinate}$ = 11). In this model, both social status and treatment were included as binomial, fixed predictors ("dominant" or "subordinate", and "control" or "stress exposure"), and an interaction between social status and treatment was included to account for possible differences in the effect of social status on mass between treatment types. Sex was included as an additional fixed predictor to control for differences in food acquisition rate that may exist between sexes, regardless of social status. Both individual identity and flight enclosure identity were initially included as random intercepts to account for variance explained by each parameter, however, flight enclosure identity was later removed because it explained negligible variance in our data (s.d. < 0.001). Finally, because previous research has suggested that the effect of stress exposure on feeding behaviour in birds can vary substantially among individuals (Favreau-Peigné et al., 2014), we allowed the variance in our model to differ according to treatment type by weighting according to binomial treatment.

We were most interested in testing whether socially subordinates experienced resource limitation across stress exposure treatments, and not control treatments. As such, we compared the change in mass experienced across stress exposure treatments alone between socially subordinate and socially dominant individuals using a planned comparison with Bonferroni correction ("emmeans" package in R; Lenth, Singmann, & Love, 2018).

#### Effect of body condition on surface temperature responses to stressors

Previous research in Blue Tits has suggested that wintering individuals with low fat reserves are more likely to display rest-phase hypothermia than those with high fat reserves, and that such hypothermia probably occurs to lessen metabolic demands of thermoregulation in the cold (Nord, Nilsson, & Nilsson, 2011). We therefore asked whether body condition alone was sufficient to explain changes in surface temperature and dry heat transfer ($q_{Tot}$) following stress exposure that occurred among socially subordinate individuals, but not socially dominant individuals. To do so, we tested whether body condition upon completion of a given treatment was influenced by both treatment type and social status, using a linear mixed effects model in the R package "glmmTMB" (Brooks et al.,

2017). Here, body condition (measured as the normalised residuals of a linear model regressing mass against wing-cord; $n$ = 69, $\beta \pm s.e.m.$ = 0.260 ± 0.035, $t$ = 7.357, $p$ < 0.001) was used as the response variable (Gaussian), whilst treatment type (factorial), social status (factorial), and an interaction between treatment and social status were included as fixed effect predictors. Sex (factorial) was also included as a fixed effect predictor to account for the possible influence of sex on condition, irrespective of social status and treatment type. Finally, individual identity (factorial) was included as a random intercept in our model to account for non-independence between samples drawn from the same individual, and residuals were weighted by treatment to correct for visually-observed heteroskedasticity in residuals between treatment types. Our final sample included 32 condition measurements drawn from 18 individuals ($n_{\text{socially dominant}}$ = 6; $n_{\text{socially subordinate}}$ = 10).

Because we were most interested in whether final body condition differed between socially dominant and socially subordinate individuals after stress exposure treatments (that is, rather than control treatments), we tested whether final condition varied across the social hierarchy following stress exposure treatment alone using a planned comparison with Bonferroni correction. Here, planned comparisons were conducted in the R package "emmeans" (Lenth et al., 2018).

#### Results

##### Social subordinates, but not social dominants, are resource limited

In our sample population, feeding rate significantly declined across time of day under control conditions ( $\beta \pm$ *s.e.m.* = -0.057 $\pm$ 0.021, $t$ = -2.712, $p$ = 0.007), but this trend was negated when individuals were exposed to repeated and rotating stressors (treatment:time of day: $\beta \pm$ *s.e.m.* = 0.063 $\pm$ 0.028, $t$ = 2.192, $p$ = 0.029; Fig. 3). Furthermore, feeding rate significantly varied across ambient temperature ($f$ = 2.192, estimated degrees of freedom, or "*e.d.f.*" = 2.786, $p$ = 0.022), with individuals displaying a bimodal response across our observed temperature range. Again, this trend was significantly altered by the presence of stress exposure ($f$ = 4.540, *e.d.f.* = 1.009, $p$ = 0.032), such that the amplitude of each mode was reduced under stress exposure conditions when compared to control conditions. Regardless of time of day and ambient temperature, treatment significantly influenced feeding rate across individuals ( $\beta \pm$ *s.e.m.* = -0.873 $\pm$ 0.321, $t$ = -2.723, $p$ = 0.007), with stress-exposed individuals globally feeding less than controls (intercept: $\beta \pm$ *s.e.m.* = 3.328 $\pm$ 0.445, $t$ = 7.483, $p$ < 0.001). Interestingly, social status did not significantly alter this suppressive effect of stress exposure on feeding (intercept: $\beta \pm$ *s.e.m.* = 0.113 $\pm$ 0.096, $t$ = 1.181, $p$ = 0.238). Social status alone, however, elicited a significant effect on feeding rate, with socially subordinate individuals feeding less than socially dominant individuals ( $\beta_{subordinate} \pm$ *s.e.m.* = -0.811 $\pm$ 0.247, $t$ = -3.287, $p$ = 0.001; SFig. 1). An effect of sex on feeding rate was not significant ( $\beta \pm$ *s.e.m.* = -0.214 $\pm$ 0.256, $t$ = -0.836, $p$ = 0.403).

In this study, we asked whether socially subordinate individuals, but not socially dominant individuals, were unable to maintain a constant body mass across stress exposure treatments. Mass did not significantly change across control treatments ($n_{days}$ = 30) for both socially subordinate ( $\beta_{\text{subordinate} \pm s.e.m.}$ = 0.032 ± 0.151, $z$ = 0.212, $p$ = 0.832) and socially dominant individuals (intercept: $\beta \pm s.e.m.$ = -0.161 ± 0.265, $z$ = -0.608, $p$ = 0.543), however, under stress exposure treatments, mass of socially subordinate individuals significantly fell (treatment:social status: $\beta \pm s.e.m.$ = -0.420 ± 0.185, $z$ = -2.270, $p$ = 0.023), whilst that of socially dominant individuals significantly increased (intercept: $\beta \pm s.e.m.$ = 0.703 ± 0.317, $z$ = 2.217, $p$ = 0.027; SFig. 2). Final change in mass across stress exposure treatments alone significantly differed between socially subordinate and socially dominant individuals, according to a planned comparison ( $\beta \pm s.e.m.$ = -0.194 ± 0.054, $t$ = -3.609, $p$ = 0.003). Similar to our results pertaining to rate of food acquisition, a global effect of sex on change in mass across treatments was not significant (sex: $\beta \pm s.e.m.$ = -0.009 ± 0.085, $z$ = -0.105, $p$ = 0.916).

#### Body condition does not significantly differ between social dominants and social subordinates on termination of stress exposure

Results of our LMM did not support a significant effect of treatment type ( $\beta \pm s.e.m.$ = 0.038 ± 0.029, $z$ = 1.321, $p$ = 0.186), social status ( $\beta \pm s.e.m.$ = 0.025 ± 0.106, $z$ = 0.231, $p$ = 0.817), or sex ( $\beta$ = 0.157 ± 0.104, $z$ = 1.505. $p$ = 0.132) on individual condition upon treatment completion. A significant interactive effect of treatment and social status was detected, however, with socially subordinate individuals alone exhibiting declines in condition across stress exposure treatments when compared to control condition ( $\beta \pm$ *s.e.m.* -0.073 $\pm$ 0.034, $z$ = -2.137. $p$ = 0.033). Despite this interactive effect, however, final condition of socially dominant and socially subordinate individuals did not significantly differ following stress exposure treatments, according to a planned comparison ($n$ = 16, $\beta \pm$ *s.e.m.* = -0.024 $\pm$ 0.053, $t$ = -0.461, $p$ = 1.000; SFig. 3).

#### Figure Captions

**Supplemental Figure 1 | Effects of social status and stress exposure on feeding rate of Black-capped Chickadees across time of day**. Feeding rate (visits/hour; marginal mean from generalised additive mixed effects model) was calculated from video observations made across 60 days, with a minimum of 1 hour of observation per flight enclosure ($n_{flight\ enclosures}$ = 4), where individuals were identified by coloured leg bands. Solid lines represent mean feeding rate of socially dominant individuals and dashed lines represent that of socially subordinate individuals. Purple and yellow bands represent 95% confidence intervals around feeding rate estimates for socially dominant and socially subordinate individuals respectively. Feeding rate significantly differed between treatments ($p$ = 0.007), and across social hierarchies ($p$ = 0.001), at $\alpha$ = 0.05.

**Supplemental Figure 2 | Effect of experimental treatment type on change in mass (across 30 days) of socially dominant and socially subordinate individuals.** Mass (g) of each individual ($n_{socially\ dominant}$ = 6; $n_{socially\ subordinate}$ = 11) was measured at the onset and completion of each treatment. Purple dots and yellow dots represent mean change in mass (marginal means from linear mixed effects model) of socially dominant and socially subordinate individuals respectively. Socially subordinate individuals lost significantly more mass that socially dominant individuals across stress exposure treatments according to planned comparison (indicated by an asterisk; $p$ = 0.023; $\alpha$ = 0.05). Whiskers represent 95% confidence intervals around means.

**Supplemental Figure 3 | Effect of experimental treatment type on condition of socially dominant and socially subordinate individuals.** Condition of each individual ($n_{socially\ dominant}$ = 6; $n_{socially\ subordinate}$ = 11) was calculated as the normalised residuals (0-100) from a linear model regression mass (g) against wing chord (mm). Purple dots and yellow dots represent mean condition (marginal means from linear mixed effects model) of socially dominant and socially subordinate individuals respectively, and whiskers represent 95% confidence intervals around means. No significant difference was detected between the condition of socially dominant and socially subordinate individuals following control or stress exposure conditions according to a linear mixed effects model and planned comparison (Control: $p$ = 0.817; stress exposure: $p$ = 1.000).

#### References


Bonter, D., Zuckerberg, B., Sedgwick, C., & Hochachka, W. (2013). Daily foraging patterns in free-living birds: Exploring the predation–starvation trade-off. *Proc R Soc B*, *280(1760)*, 20123087.

Brittingham, M., & Temple, S. (1992). Use of winter bird feeders by black-capped chickadees. *J Wildl Manag*, *55(1)*, 103–111.

Brooks, M., Kristensen, K., van Benthem, K., Magnusson, A., Berg, C., Skaug, A. N. H., … Bolker, B. (2017). GlmmTMB balances speed and flexibility among packages for zero-inflated generalized linear mixed modeling. *R Journal*, *9(2)*, 378–400. Retrieved from https://journal.r-project.org/archive/2017/RJ-2017-066/index.htm

Evans, J., Devost, I., Jones, T., & Morand-Ferron, J. (2018). Inferring dominance interactions from automatically recorded temporal data. *Ethol*, *124(3)*, 188–195.

Favreau-Peigné, A., Calandreau, L., Constantin, P., Gaultier, B., Bertin, A., Arnould, C., … Boissy, A. (2014). Emotionality modulates the effect of chronic stress on feeding behaviour in birds. *PLoS One*, *9(2)*, e87249.

Knauer, J., El-Madany, T., Zaehle, S., & Migliavacca, M. (2018). Bigleaf - an r package for the calculation of physical and physiological ecosystem properties from eddy covariance data. *PLoS ONE*, *13*(8), e0201114. doi: 10.1371/journal.pone.0201114

Lenth, R., Singmann, H., & Love, J. (2018). Emmeans: Estimated marginal means, aka least-squares means. *R Package Version*, *1(1)*.

Neumann, C., & Kulik, L. (2019). EloRating: Animal dominance hierarchies by Elo rating. *R Package, Version 0.46.8*.

Nord, A., Nilsson, J., & Nilsson, J. (2011). Nocturnal body temperature in wintering blue tits is affected by roost-site temperature and body reserves. *Oecologia*, *167(1)*, 21–25.

R Core Team. (2019). R: A language and environment for statistical computing. r foundation for statistical computing, vienna, austria. *Https://www.R-Project.org/*.

Reinertsen, R., & Haftorn, S. (1984). The effect of short-time fasting on metabolism and nocturnal hypothermia in the willow tit, *parus montanus*. *J Comp Physiol B*, *154(1)*, 23–28.

Robertson, J., Mastromonaco, G., & Burness, G. (2020). Evidence that stress-induced changes in surface temperature serve a thermoregulatory function. *J Exp Biol*, *223(4)*, jeb213421.

Silva, L., Lardy, S., Ferreira, A., Rey, B., Doutrelant, C., & Covas, R. (2018). Females pay the oxidative cost of dominance in a highly social bird. *Anim Behav*, *114(1)*, 135–146.

Snijders, L., Naguib, M., & van Oers, K. (2016). Dominance rank and boldness predict social attraction in great tits. *Behav Ecol*, *28(2)*, 398–406.

Tattersall, G. (2019). *Thermimage: Thermal image analysis*. doi: 10.5281/zenodo.1069704

Wood, S. (2011). Fast stable restricted maximum likelihood and marginal likelihood estimation of semiparametric generalized linear models. *J R Stat Soc B*, *73(1)*, 3–36.
